# Sex-specific regulation of social play in juvenile rats by oxytocin neurons in the hypothalamus and oxytocin signaling in the nucleus accumbens

**DOI:** 10.1101/2025.11.11.687699

**Authors:** Samantha M. Bowden, Kira D. Becker, Anna Luxhoj, Valery Grinevich, Jessica DA Lee, Alexa H. Veenema

## Abstract

Social play is a rewarding behavior shown across juvenile mammalian species and is important for the development of social competency throughout the lifespan. The neuropeptide oxytocin (OXT) regulates various social behaviors and is being used in clinical trials to improve social competency. However, the role of OXT in juvenile social play is largely unknown. To address this gap, we determined the involvement of hypothalamic OXT-producing neurons in the supraoptic nucleus (SON) and paraventricular nucleus (PVN), PVN^OXT^ projections to the nucleus accumbens (NAc), and OXT signaling within the NAc, in the regulation of social play in juvenile male and female rats. We found that neither chemogenetic stimulation of SON^OXT^ cell bodies nor PVN^OXT^ projections to the NAc altered the expression of juvenile social play but instead increased social investigation. However, chemogenetic stimulation of PVN^OXT^ cell bodies as well as intra-NAc infusion of OXT decreased social play in males without an effect in females. Lastly, social play duration correlated negatively with the proportion of activated NAc^OXTR^ neurons, an effect driven by males. Together, these findings suggest that distinct OXT neuronal populations modulate different forms of social behavior and that PVN^OXT^ neurons and NAc-OXT signaling sex-specifically modulate social play behavior.

## INTRODUCTION

Social play (a.k.a rough and tumble play or play fighting) is a highly rewarding behavior (Trezza et al., 2010; Vanderschuren et al., 2016) displayed by many mammalian species, including rats, non-human primates, and humans (Calcagnetti & Schechter, 1992; Panksepp, 1981; Anthony D. Pellegrini & Smith, 2005; Pellis & Pellis, 1997; Shimada & Sueur, 2018). Social play peaks during the juvenile period and is necessary for the expression of context-appropriate social behaviors through life (Anthony D. Pellegrini & Smith, 1998; A D Pellegrini & Smith, 1998; A D Pellegrini, 1989; van den Berg et al., 1999). Children with social disorders such as autism spectrum disorder (ASD show decreased participation in social play which is thought to contribute to their reduced ability to appropriately navigate social and emotional situations (Chevallier et al., 2012, 2013; Jordan, 2003).

The neuropeptide oxytocin (OXT) is being tested for its potential to improve social engagement in ASD adults and children (clinicaltrials.gov). This is largely based on decades of research in adult rodents implicating OXT in the regulation of various social behaviors (Oettl et al., 2016; Olazábal & Young, 2006a; Heather E Ross et al., 2009). OXT is primarily produced in the paraventricular nucleus of the hypothalamus (PVN) and supraoptic nucleus (SON; (Sawchenko & Swanson, 1983; Tang et al., 2020). PVN^OXT^ neurons in particular are involved in the regulation of social behaviors (Grund et al., 2019; Hung et al., 2017; Oettl et al., 2016; Tang et al., 2020). For example, PVN^OXT^ neuronal activity increased in paternal male mandarin voles when they were licking and retrieving their pups (He et al., 2021). Furthermore, PVN^OXT^ neuronal activity increased in adult male mice while investigating a juvenile mouse (Hung et al., 2017). In the same study, optogenetic stimulation of PVN^OXT^ neurons increased juvenile investigation in adult male mice (Hung et al., 2017). Moreover, optogenetic inhibition of PVN^OXT^ neurons decreased social interaction among adult female rats (Tang et al., 2020). The involvement of SON^OXT^ neurons in social behavior is less well studied, with thus far one study showing that chemogenetic inhibition of SON^OXT^ neurons projecting to the lateral septum (LS) decreased investigation of an adult female mouse in lactating female mice (Menon et al., 2018). These findings suggest that activation of PVN^OXT^ and SON^OXT^ neurons facilitates the expression of social behaviors in adult rodents. However, it is unclear how these two populations of OXT neurons modulate social behaviors in juvenile rats.

PVN^OXT^ neurons project to several regions in the brain, including those regulating rewarding behaviors, such as the nucleus accumbens (NAc; (Knobloch et al., 2012; H E Ross et al., 2009; Tang et al., 2020). Chemogenetic inhibition of PVN^OXT^ terminals in the NAc decreased pup care and increased the latency to retrieve pups in male mandarin voles (He et al., 2021). Additionally, optogenetic stimulation of PVN^OXT^ fibers into the NAc in both male and female mandarin voles increased their time investigating a sex- and age-matched conspecific (Hou et al., 2023). These findings suggest that activation of the PVN^OXT^ to NAc pathway facilitates the expression of social behaviors in adult rodents, but whether this is also true for juvenile rodents is unknown.

The NAc is an OXT receptor (OXTR) dense region in both juvenile and adult rats (Klawonn & Malenka, 2018; Mitre et al., 2016; Smith et al., 2017). OXT signaling in the NAc is involved in the regulation of social behaviors in adult rodents (Insel & Shapiro, 1992; Liu & Wang, 2003; Olazábal & Young, 2006a; Yu et al., 2016). For example, OXTR binding density in the NAc correlates positively with the level of maternal care and with partner preference formation in female prairie voles (Insel & Shapiro, 1992; Olazábal & Young, 2006a). Additionally, overexpression of OXTRs in the NAc facilitates and OXTR antagonism in the NAc blocks the formation of a partner preference (Keebaugh & Young, 2011; Young et al., 2001), suggesting that OXT signaling in the NAc is both sufficient and necessary. Furthermore, OXTR antagonism blocks the formation of a conditioned place preference for a social stimulus in adolescent male mice (Dölen et al., 2013),suggesting OXT signaling in the NAc may mediate the perceived pleasurable aspect of social interactions. However, it is yet to be determined whether OXT signaling in the NAc mediates the expression of juvenile-typical social behaviors, such as social play.

The goal of the present study was to determine the involvement of PVN^OXT^ and SON^OXT^ neurons, the PVN^OXT^ to NAc pathway, and OXTergic signaling within the NAc in the regulation of social play in juvenile male and female rats. We hypothesized that increased activation of OXT neurons in the PVN and SON, increased activation of the PVN^OXT^ to NAc pathway, and increased OXT signaling in the NAc promote the expression of social play behavior in both sexes. To test this, we first determined the effects of chemogenetic stimulation of PVN^OXT^ neurons and SON^OXT^ neurons on the expression of social play behaviors (Experiments 1 and 2). Next, we determined whether PVN^OXT^ neurons project to the NAc and whether these projections are similar across sex (Experiment 3). Based on these outcomes, we then determined the effects of chemogenetic stimulation of PVN^OXT^ terminals within the NAc on the expression of social play behaviors (Experiment 4). We also examined the effects of synthetic OXT infusion into the NAc on the expression of social play behaviors (Experiment 5). Finally, we used *in situ* hybridization to determine whether social play exposure alters the activation of *oxtr* mRNA-expressing neurons in the NAc (Experiment 6).

## METHODS

### Animals

For experiments 1-4, male and female Wistar rats were bred in-house using animal stock previously obtained from Charles River Laboratories (Raleigh, NC, USA). Experimental rats were weaned at 21 days of age and housed in single-sex groups of two in standard Allentown cages (48x27x20cm). For experiment 5, 25-day-old male and female Wistar rats were obtained from Charles River Laboratories (Raleigh, NC, USA), where they were housed in the same condition as in-house bred rats following arrival. For all experiments, stimulus male and female Wistar rats used for play testing were obtained from Charles River Laboratories (Raleigh, NC, USA; 26-30 days of age at arrival) and were housed in same-sex groups of three to four for the duration of the experiment. All experimental and stimulus rats were maintained under standard laboratory conditions (12:12h light/dark cycle, lights off at 13:00h, 22°C, 50% humidity, food, and water available *ad libitum*). All experiments were conducted in accordance with the National Institute of Health *Guidelines for Care and Use of Laboratory Animals* and approved by the Michigan State University Institutional Animal Care and Use Committee.

### Social Play Testing

Rats were tested for social play in Experiments 1, 2, 4, and 5. Social play testing occurred between 31-35 days of age beginning at lights out (13:00h). Experimental rats were singly housed in a clean cage twenty-four hours prior to the start of social play testing. During the test, a novel sex- and age-matched conspecific was introduced into the experimental rats’ home cage and they were allowed to freely interact for 10 min. Play sessions were video recorded and behavior of the experimental rat was later scored using the video analysis program Solomon Coder (Solomon.andraspeter.com) by a researcher blind to the experimental conditions. Duration of social play (time spent engaging in playful interactions with the stimulus rat including chasing, nape attacks, wrestling, and pinning), social investigation (sniffing the anogenital region of the stimulus rat), allogrooming (grooming head and neck of stimulus rat), total duration of social behavior (cumulative percentage of time spent engaging in social play, social investigation and allogrooming), and cage exploration, along with number of pins (experimental rat holds stimulus rat in supine position), nape attacks (experimental rat attacks or makes nose contact with the nape of the neck of the stimulus rat), and supine positions (experimental rat rolls on its back or is pinned on its back by stimulus rat) were scored according to (Bredewold et al., 2018).

### Fluorescent Immunohistochemistry (FIHC) and Image acquisition

At the conclusion of each experiment, rats were deeply anesthetized with isoflurane (5%) before transcardial perfusion with 0.9% saline followed by 4% paraformaldehyde in 0.1 M borate buffer (pH= 9.5). Brains were extracted and post-fixed in 4% paraformaldehyde with 12% sucrose for 24 h. The olfactory bulb and cerebellum were removed using a brain matrix. Brains were flash frozen in 2-methlybutane and stored at -80°C until histological processing. Brains were sliced on a cryostat (Leica CM3050, Buffalo Grove, IL) in 30 µm coronal sections into four consecutive series containing region(s) of interest. Brain sections were stored at -20°C in cryoprotectant solution (0.05 M sodium phosphate buffer, 30% ethylene glycol, 20% glycerol) until undergoing FIHC staining.

For FIHC processing, sections were washed in tris-buffered saline (TBS; 0.1 M; pH=7.4) and incubated overnight at 4°C in a primary antibody solution (See Table 1 for full antibody information) with 2 % normal donkey serum as a blocking agent (Jackson ImmunoResearch, West Grove, PA) and 0.3 % triton-X (Sigma-Aldrich, Burlington, VA). The following day, tissue was washed in TBS and incubated in a fluorescent secondary antibody solution at room temperature for 1 h (See Table 1 for full antibody information). After rinsing in TBS, tissue was mounted onto Superfrost slides (Fisher Scientific, Waltham, MA, USA), air-dried, and coverslipped with a Vectashield hardset antifade mounting medium with a DAPI counterstain (Vector Laboratories, Burlingame, CA).

All images of FIHC processed tissue were acquired with a 20x objective on a Keyence BZ-X700E/BZ-X710 fluorescent microscope and associated BZ-H3AE software (Keyence Corporation of America, Elmwood Park, NJ).

**Table 1:** FIHC antibodies.

| Antibody Name | Clonality/<br>Species | Company | Product No. | Dilution |
| --- | --- | --- | --- | --- |
| Anti-mCherry | Chicken<br>monoclonal | Abcam | ab205402 | 1:2000 |
| Anti-GFP | Chicken<br>monoclonal | Invitrogen | A10262 | 1:2000 |
| Anti-OXT | Mouse<br>monoclonal | Gift from Hal<br>Gainer | NA | 1:5000 |
| AlexaFluor 488 | Donkey anti-<br>chicken | Jackson<br>Laboratories | 703-545-155 | 1:500 |
| AlexaFluor 488 | Donkey anti-<br>mouse | Jackson<br>Laboratories | 715-545-150 | 1:500 |
| AlexaFluor 594 | Donkey anti-<br>chicken | Jackson<br>Laboratories | 703-585-155 | 1:250-1:500 |

### Experiment 1: Effects of chemogenetic stimulation of PVN^OXT^ neurons on social play behavior

To determine the role of PVN^OXT^ neurons in the regulation of juvenile social play, we chemogenetically stimulated OXT neurons in the PVN. At 26 days of age, male and female rats underwent stereotaxic surgery in which an excitatory designer receptor exclusively activated by designer drugs (DREADD) viral vector (AAV1/2-OXTp-hM3Dq-mCherry; n=7 males and n=9 females) or mCherry control viral vector (AAV1/2-OXTp-mCherry; n=6 males and n=7 females) under the control of the murine OXT promoter was infused bilaterally into the PVN (200 nl/hemisphere). Rats were anesthetized with isoflurane (2-4% as needed; Henry Schein, Melville, NY, USA) and mounted onto a stereotaxic frame (Stoelting, Wood Dale, IL, USA). A 1 ml 7000 series Hamilton syringe (Hamilton, Reno, NV, USA) was attached to a stereotaxic injector system (Stoelting, Wood Dale, IL, USA) and loaded with AAV1/2-OXTp-hM3Dq-mCherry or AAV1/2-OXTp-mCherry. The syringe was aimed at the PVN (AP: -1.70 mm ML: ±1.70 mm DV: -7.70 mm from bregma in accordance with the Rat Brain Atlas; (Paxinos & Watson, 2013) at a 10° angle to avoid damage to the sagittal sinus. The viral vector was infused at a rate of 100 nl/min, and the needle remained in place for 10 min to allow adequate time for diffusion and uptake of the viral vector. The syringe was slowly withdrawn, and the process was repeated in the contralateral PVN. Rats received a subcutaneous injection of meloxicam (2 mg/kg; Covetrus) following infusion and once a day for two days after surgery. Rats were singly housed and monitored post-surgery until fully ambulatory, after which they were pair-housed with same-sex littermates that underwent the same stereotaxic surgery until 24 hours prior to the onset of behavioral testing.

Beginning the day after surgery, experimental rats were habituated to the intraperitoneal (IP) injection procedure. Once a day for three days, experimental rats were removed from their homecage, held by their scruff, and an empty syringe was inserted into the abdomen at a 90° angle. Play testing occurred over two days (at 33 and 35 days of age) with a rest day in between trials (see Figure 1 for timeline). IP injections of either clozapine-*N-*oxide (CNO; 0.3 mg/kg) or sterile 0.9% saline (0.02 ml) were administered 30 min prior to testing in a randomly assigned counterbalanced manner.

Sections containing the PVN (Bregma -1.08 mm to -1.72 mm; Paxinos and Watson, 2013) were processed for mCherry cell body detection using FIHC. Images of the mCherry FIHC were acquired to verify the location of DREADD transfection. Rats with bilateral mCherry expression in the PVN were included in subsequent behavioral analysis.

### Experiment 2: Effects of chemogenetic stimulation of SON^OXT^ neurons on social play behavior

To determine the role of SON^OXT^ neurons in the regulation of juvenile social play, we chemogenetically stimulated OXT neurons in the SON. At 26 days of age, male and female rats underwent stereotaxic surgery in which an excitatory DREADD viral vector under the control of the murine OXT promoter (AAV1/2-OXTp-hM3Dq-mCherry; n=6/sex) was infused bilaterally into the SON. All surgical, behavioral, FIHC, and imaging procedures were performed as described in experiment 1 with the following exceptions: The syringe was aimed at the SON (AP: -1.30mm ML: ±1.65mm DV: -8.50mm from bregma in accordance with the Rat Brain Atlas; Paxinos and Watson, 2013), and sections containing the SON (Bregma -0.36 to -1.44 mm; Paxinos and Watson, 2013) were processed for mCherry cell body detection. Rats with bilateral mCherry expression in the SON were included in subsequent behavioral analysis.

### Experiment 3: Characterization of PVN^OXT^ fibers projecting to the NAc

To better understand the role of OXT signaling in the regulation of juvenile social play behavior, we focused on downstream brain regions of the PVN. We selected the NAc, because it receives OXTergic input from the PVN (Tang et al., 2020), is an OXTR dense node (Insel & Shapiro, 1992; Smith et al., 2019) and is part of the mesolimbic reward pathway (Floresco, 2015; Salgado & Kaplitt, 2015). We first determined the PVN^OXT^ innervation pattern within the NAc of juvenile male and female rats. In detail, we infused a mCherry or Venus fluorescent reporter virus under the control of the OXT promoter (AAV1/2-OXTp-mCherry or AAV1/2-OXTp-Venus) into the PVN of 22-day-old male and female rats as described in experiment 1 (n=4 males and n=6 females). Rats were singly housed and monitored post-surgery until fully ambulatory, after which they were housed in same-sex groups of two or three until transcardial perfusion and brain extraction at 35 days of age.

30 µm coronal sections containing the NAc (+3.24 mm to +1.08 mm relative to bregma; Paxinos and Watson, 2013) and the PVN (-1.08 mm to -1.72 mm relative to bregma; Paxinos and Watson, 2013) were each collected in four consecutive series. One PVN series was processed to detect mCherry or Venus using FIHC.

Images were taken of the anterior NAc shell+core, intermediate NAc shell, intermediate NAc core, posterior NAc shell, and posterior NAc core (Figure 4B; Bregma +2.76mm to +2.28mm, +1.80mm to +1.56mm, and +1.08mm to +0.84mm, respectively; Paxinos and Watson, 2013). Images were converted into 8-bit in ImageJ (NIH; imagej.nih.gov/ij) and the number of pixels representing mCherry or Venus-positive fibers was calculated using the integrated density (IntDen) function. An average background density (BGD) measurement was acquired per image, and adjusted integrated density was calculated as (IntDen – (Area of Image*Average BGD)).

### Experiment 4: Effects of chemogenetic stimulation of PVN^OXT^ terminals in the NAc on social play behavior

Following the identification of the anterior NAc as the subregion receiving the highest density of PVN^OXT^ innervation, we next aimed to determine the role of the PVN^OXT^ to NAc pathway in the regulation of juvenile social play behavior. To accomplish this, we chemogenetically stimulated PVN^OXT^ terminals within the anterior NAc. To this end, an AAV1/2-OXTp-hM3Dq-mCherry (200nl/hemisphere) was infused into the PVN of 22-day-old male and female Wistar rats (n=7/sex) following surgical procedures as described in experiment 1. At 28-29 days of age, rats underwent a second stereotaxic surgery for bilateral implantation of 6 mm guide cannulae 2 mm dorsal to the NAc (AP: -2.6 mm, ML: +/-2.5 mm, DV: -3.25 mm from Bregma, Paxinos and Watson, 2013). This allowed for administration of CNO into the NAc to stimulate the PVN^OXT^ terminal fibers in the NAc. All rats were pair-housed with same-sex littermates that underwent stereotaxic surgery until 24 hours prior to the start of behavioral testing.

Beginning the day after cannula implantation, experimental rats were habituated to the handling and microinjection procedure once a day for three days in which the rats were wrapped tightly in a towel followed by removal of the dummy cannula and placement of the internal cannula in the guide that was left in place for 30s after which the internal canula was removed, the dummy canula was replaced and the rats was released in its homecage. Social play testing occurred over two days (at 33 and 35 days of age) with a rest day in between trials (see Figure 5A for experimental timeline). Twenty min prior to social play testing, experimental rats received bilateral infusions of either sterile 0.9% saline or 1 mM CNO into the NAc. The infusions (0.3 μL/hemisphere) were given over the course of 45 sec via an internal cannula (28 gauge; Plastics One, Roanoke, VA) that extends 2 mm beyond the guide cannula and was connected via polyethylene tubing to a 2 μL syringe (Hamilton Company #88400) mounted onto a microinfusion pump (GenieTouch, Kent Scientific, Torrington, CT). The internal cannula was kept in place for an additional 30s following infusion to allow for tissue uptake before being replaced by the dummy cannula. Drug conditions (CNO, saline) were randomly assigned in a counterbalanced manner over the two days of testing.

At the conclusion of the second day of testing, experimental rats were perfused and brains were collected for histological processing as described in experiment 3 with the following exceptions: One series of the NAc was mounted onto gelatin-subbed slides and counterstained with thionin to determine cannula placement location. One series of the PVN was processed for mCherry-ir cells via FIHC to confirm viral transfection. Rats with both bilateral cannula placement in the NAc and bilateral mCherry staining contained within in the PVN were used for behavioral analysis.

### Experiment 5: Effects of acute OXT infusion into the NAc on social play behavior

We determined the effects of an acute OXT infusion into the NAc on the expression of juvenile social play behavior. At 28 days of age, male and female Wistar rats were anesthetized with isoflurane (2-4% as needed; Henry Schein, Melville, NY, USA) and mounted onto a stereotaxic frame (Stoelting, Wood Dale, IL, USA) for bilateral implantation of 6 mm guide cannulae 2 mm dorsal to the NAc (AP:-2.6mm, ML:+/-2.5mm, DV:-3.25mm from Bregma, Paxinos and Watson, 2013; n=6 males and n=5 females). All surgical procedures, infusion habituation, and social play testing occurred as described in experiment 4 with the exception that 20 mins prior to social play testing experimental rats received bilateral infusions of either sterile 0.9% saline or OXT (1µg/0.3µl [low dose], 5µg/0.3µl [high dose]) into the NAc. The dose of OXT was randomly assigned in a counterbalanced manner, with all experimental rats receiving the three drug treatments over three days of testing.

At the conclusion of the third day of testing, rats were euthanized with CO_2_, and charcoal was injected as a marker to check proper placement of the cannulae into the NAc. Brains were then extracted, flash frozen in 2-methlybutane, and stored at -80°C until histological processing. The NAc was sectioned at 16µm onto Superfrost Plus slides (Fisher Scientific, Waltham, MA, USA) using a cryostat (Leica CM3050, Buffalo Grove, IL). Slides were counterstained with thionin to determine cannula placement. Rats with bilateral charcoal placement within the NAc were used for subsequent behavioral analysis.

### Experiment C: Activation patterns of NAc^OXTR^ expressing neurons in response to social play

We aimed to determine how social play alters activation of NAc^OXTR^ expressing neurons in juvenile male and female rats and whether activation patterns differ by sex. All rats were housed in same-sex pairs until 33 days of age, at which time rats were assigned to either “No Social Play” (n=8/sex) or “Social Play” (n=8 males and n=6 females) condition and single-housed in clean cages. The pair-housed cages of the rats in the “Social Play” condition were used as the social play testing environment. Beginning at lights out the following day, experimental rats in the “Social Play” condition were rejoined with their previous cagemate in their original pair-housed cage for the 10-minute social play test. One randomly selected rat in each play pair was striped with a permanent marker 30-60 minutes prior to the social play text to distinguish the two rats during later video analysis. After the 10-minute test, rats were returned to their single-housed cages. Concurrently, cages containing rats in the “No Social Play” condition were removed from the housing rack and placed on a table for 10 minutes to mimic the environment experienced by rats in the “Social Play” condition. Thirty-five minutes after the start of the social play test (“Social Play” condition) or removal from the housing rack (“No Social Play” condition), all rats were transcardically perfused, and brains were extracted.

30 µm coronal sections containing the NAc (+3.24 mm to +1.08 mm relative to bregma; Paxinos and Watson, 2013) were collected in four consecutive series. One series of the NAc was processed for fluorescent *in situ* hybridization to visualize *oxtr* and *fos* mRNA-expressing neurons. Blocked tissue sections containing only the NAc were mounted onto Superfrost Plus slides (Fisher Scientific). RNAScope™ Multiplex Fluorescent Reagent V2 Kits (323100, Advanced Cell Diagnostics) and probes to detect *oxtr* and *fos* mRNA were used according to user manual instructions from the supplier (Document Number 323100-USM, Advanced Cell Diagnostics). Briefly, tissue sections were washed in a phosphate buffer solution (pH: 7.6), dried at 60°C (30 min), then post-fixed in 4% paraformaldehyde (15 min) followed by dehydration in an ethanol series. Following hydrogen peroxide incubation (10 min) and target retrieval in a steamer at 99°C (5 min), tissue was then treated with protease III (30 min; 322340, Advanced Cell Diagnostics) at room temperature (25 min). The fos-C1 (403591, Advanced Cell Diagnostics) and oxtr-C3 probes were then hybridized in a HybEZTM oven (2 h; Advanced Cell Diagnostics) at 40°C. After probe hybridization, tissue sections were incubated with amplifier probes (AMP1, 40°C, 30 min; AMP2, 40°C, 30 min; AMP3, 40°C, 15 min). *fos* mRNA was tagged to the fluorophore fluorescein (1:1500; NEL741E001KT, Akoya Biosciences, 40°C, 30 min) and *oxtr* mRNA was tagged to the fluorophore Cyanine5 (Cy5; 1:1500; NEL705A001KT, Akoya Biosciences, 40°C, 30 min). Slides were then rinsed in wash buffer and coverslipped with Vectashield hardset antifade mounting medium with a DAPI counterstain (H-1500-10, Vector Laboratories) and stored at 4°C. All images of the NAc were acquired with a 20x objective on a Keyence BZ-X700E/BZ-X710 fluorescent microscope and associated BZ-H3AE software (Keyence Corporation of America, Elmwood Park, NJ). Cells were counted as *fos*+ if they had a halo-like puncta pattern around the nucleus and cells were counted as *oxtr+* if they had 3 or more puncta. In each image, the total number of *fos+, oxtr+,* and *fos+oxtr+* double-labeled neurons were counted by an experimenter blind to sex and social play condition. In addition to raw counts, the proportion of *oxtr+* neurons that were *fos+* ([*oxtr+fos+/fos+*]x100) and the proportion of *fos+* neurons that were *oxtr+* ([*oxtr+fos+/oxtr+*]x100) were manually calculated for each rat.

### Statistical Analysis

For experiments 1, 2, 4, and 5, the effects of sex and drug on behavior were analyzed using a 2-way within subject analysis of variance (ANOVA), with experiment 5 using a Greenhouse-Geisser correction for violations of sphericity. For experiment 3, effects of sex and NAc subregion (Anterior NAc, Intermediate Core, Intermediate Shell, Posterior Core, Posterior Shell) in PVN^OXT^ fiber density in the NAc were assessed using a 2-way mixed-model ANOVA. For experiment 6, the effects of sex and social play condition on NAc^OXTR^ activation patterns were assessed using a 2-way between subjects ANOVA, and sex differences in social play behaviors were assessed using Student’s unpaired t-tests. Given baseline sex differences in social play in experiment 6, correlation coefficients were assessed using Pearson’s R for each group and separated by sex when a significant correlation occurred. Holm-Sidak post-hoc tests were performed when significant main effects (experiments 3 and 5) or interactions (all experiments) occurred. Effect sizes were calculated using partial η^2^ (η^2^_p_=SS_effect_/SS_residual_ for repeated measures and mixed effect ANOVAs. Analyses were performed in GraphPad Prism (version 10.6) significance was set at *p* < 0.05.

## RESULTS

### Experiment 1: Chemogenetic stimulation of PVN^OXT^ neurons decreases social play behaviors in juvenile male rats

There was a significant sex x drug interaction on the duration of social play, the number of nape attacks, and the number of pins (See Table 2 for statistical details). Post-hoc Holm-Sidak’s test for multiple comparisons revealed that systemic administration of CNO significantly reduced the duration of social play (*p* = 0.006; Figure 1G) and the number of nape attacks (*p* = 0.048; Figure 1H) in males, while a non-significant trend towards an increase in the duration of social play was observed in females (*p =* 0.059; Figure 1G). Additionally, there was a trend towards a baseline sex difference in social play duration with saline treated males showing longer duration of social play compared to saline treated females (*p =* 0.051), which was eliminated following CNO treatment (*p* = 0.88; Figure 1G). Post-hoc analysis did not reach significance regarding the number of pins for any condition. There was no effect of sex, drug, or sex x drug interactions on the duration of social investigation, duration of allogrooming, or cumulative duration of social behaviors (*p* > 0.05 for all, Figure 1J-L; see Table 2.1 for full analysis). Lastly, there was no effect of systemic CNO administration in the absence of DREADDs (OXTp-mCherry controls) on any behaviors quantified, indicating that CNO did not alter social behavior in the absence of hM3Dq (*p* > 0.05 for all, Supplemental Figure 2A-F; see Supplemental Table 1 for statistical details).

**Figure 1.**
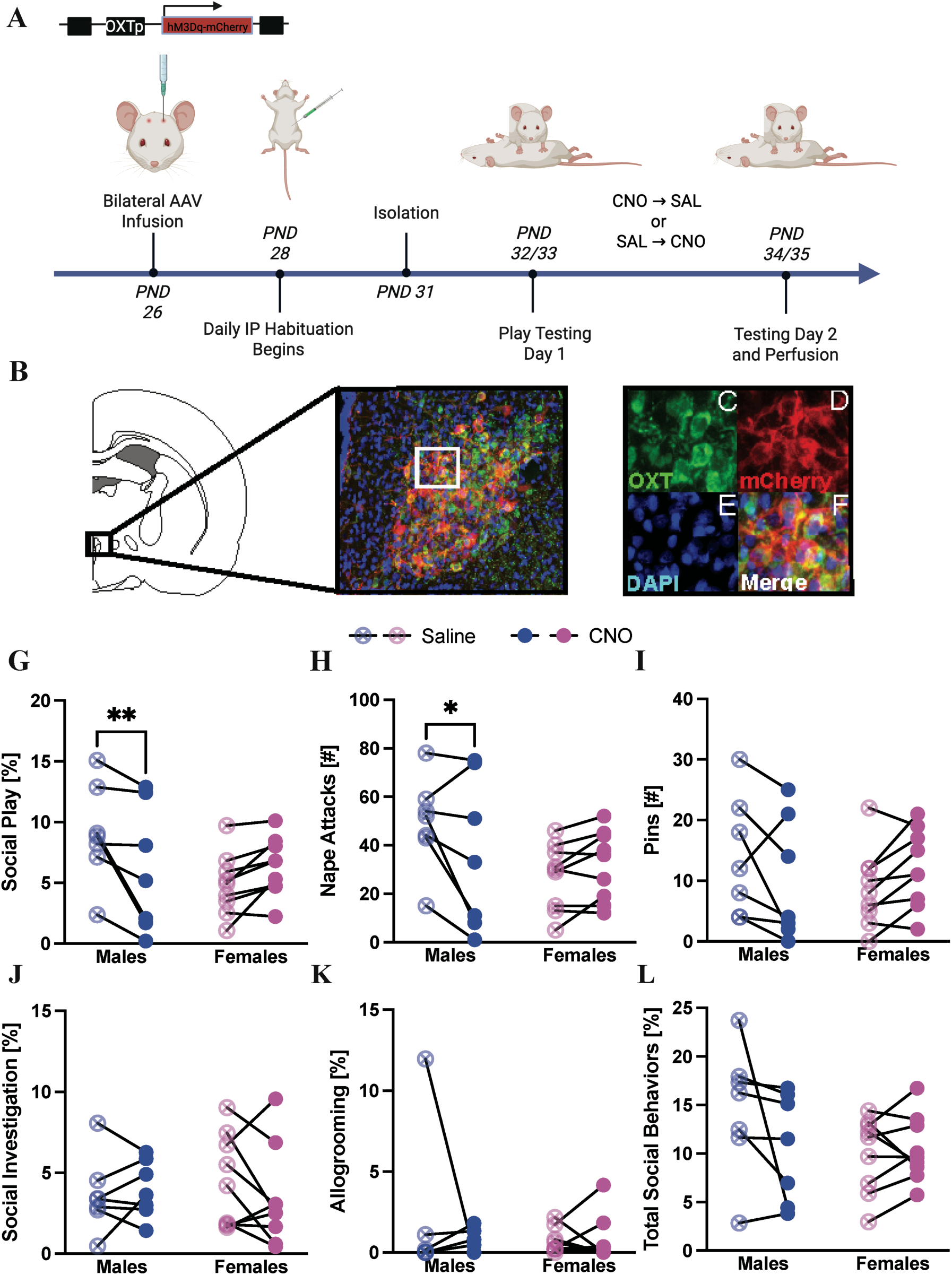
Chemogenetic stimulation of PVN^OXT^ neurons alters social play behaviors sex-specifically in juvenile rats. (A) Timeline of experimental procedures in which juvenile rats (26-day-old) underwent stereotaxic surgery for infusion of AAV-OXTp-hM3Dq-mCherry into the PVN. Following recovery and habituation to IP injections, all rats underwent two days of social play testing in which either saline or CNO was administered 30 min prior to testing. Drug conditions were randomly assigned and counterbalanced over the two days of testing. (B) Rat brain atlas image (modified from Paxinos and Watson, 2007) indicating the location and representative image of mCherry expression within PVN^OXT^ neurons. (C-F) Magnifications of image in B denoting mCherry (C, red) and OXT (D, green) positive neurons that are co-localized (F, yellow) using DAPI (E) as a cellular marker. (G-L) Administration of CNO significantly decreased the duration of social play in males and had a strong tendency towards increasing the duration of social play in females (G) as well as significantly decreased the number of nape attacks in males (H). CNO administration did not alter number of pins (I), duration of social investigation (J), duration of allogrooming (K), or total duration of social behaviors (L). Social play, investigation, allogrooming and combined social behaviors are expressed as a percentage of total time. 2-way ANOVA, Sex x Drug interaction; *: *p* < 0.05, **: *p* < 0.01; Holm’s-Sidak post-hoc tests.

**Table 2.** Experiment 1: Effects of chemogenetic stimulation of PVN^OXT^ neurons; Two-way ANOVA statistics for behaviors quantified in social play testing. Significant main effects and interactions are notated in **bold**.

|  | Drug Treatment | Sex | Drug x Sex |
| --- | --- | --- | --- |
| Social Play [%] | $F_{1,14} = 1.85$,<br>$p = 0.20$, $\eta^2_p = 0.12$ | $F_{1,14} = 1.33$,<br>$p = 0.27$, $\eta^2_p = 0.48$ | **$F_{1,14} = 16.7$,**<br>**$p = 0.001$, $\eta^2_p = 0.54$** |
| Nape Attacks [#] | $F_{1,14} = 1.54$,<br>$p = 0.24$, $\eta^2_p = 0.10$ | $F_{1,14} = 1.92$,<br>$p = 0.19$, $\eta^2_p = 0.50$ | **$F_{1,14} = 6.52$,**<br>**$p = 0.02$, $\eta^2_p = 0.32$** |
| Pins [#] | $F_{1,14} = 0.06$,<br>$p = 0.81$, $\eta^2_p = 0.004$ | $F_{1,14} = 0.16$,<br>$p = 0.69$, $\eta^2_p = 0.07$ | **$F_{1,14} = 6.73$,**<br>**$p = 0.02$, $\eta^2_p = 0.33$** |
| Social Investigation [%] | $F_{1,14} = 0.44$,<br>$p = 0.52$, $\eta^2_p = 0.03$ | $F_{1,14} < 0.01$,<br>$p = 0.93$, $\eta^2_p = 0.002$ | $F_{1,14} = 1.63$,<br>$p = 0.22$, $\eta^2_p = 0.11$ |
| Allogrooming [%] | $F_{1,14} = 0.54$,<br>$p = 0.47$, $\eta^2_p = 0.04$ | $F_{1,14} = 0.48$,<br>$p = 0.50$, $\eta^2_p = 0.03$ | $F_{1,14} = 0.63$,<br>$p = 0.44$, $\eta^2_p = 0.04$ |
| Total Social Behavior [%] | $F_{1,14} = 1.53$,<br>$p = 0.24$, $\eta^2_p = 0.11$ | $F_{1,14} = 1.58$,<br>$p = 0.23$, $\eta^2_p = 0.20$ | $F_{1,14} = 3.23$,<br>$p = 0.09$, $\eta^2_p = 0.17$ |

### Experiment 2: Chemogenetic stimulation of SON^OXT^ neurons decreases the number of pins and increases social investigation in juvenile male and female rats

There were significant main effects of drug condition on the number of pins and the duration of social investigation, with CNO administration decreasing the number of pins (Figure 2I) and increasing the duration of social investigation (Figure 2J) compared to saline administration (See Table 3 for statistical details). Furthermore, there were significant main effects of sex on the duration of social play and the number of nape attacks. Here, females showed a shorter duration of social play and fewer nape attacks (Figure 2H) than males. There were no significant main effects or interaction effects on allogrooming or total duration of social behavior (*p* > 0.05 for both, Figure 2K-L).

**Figure 2.**
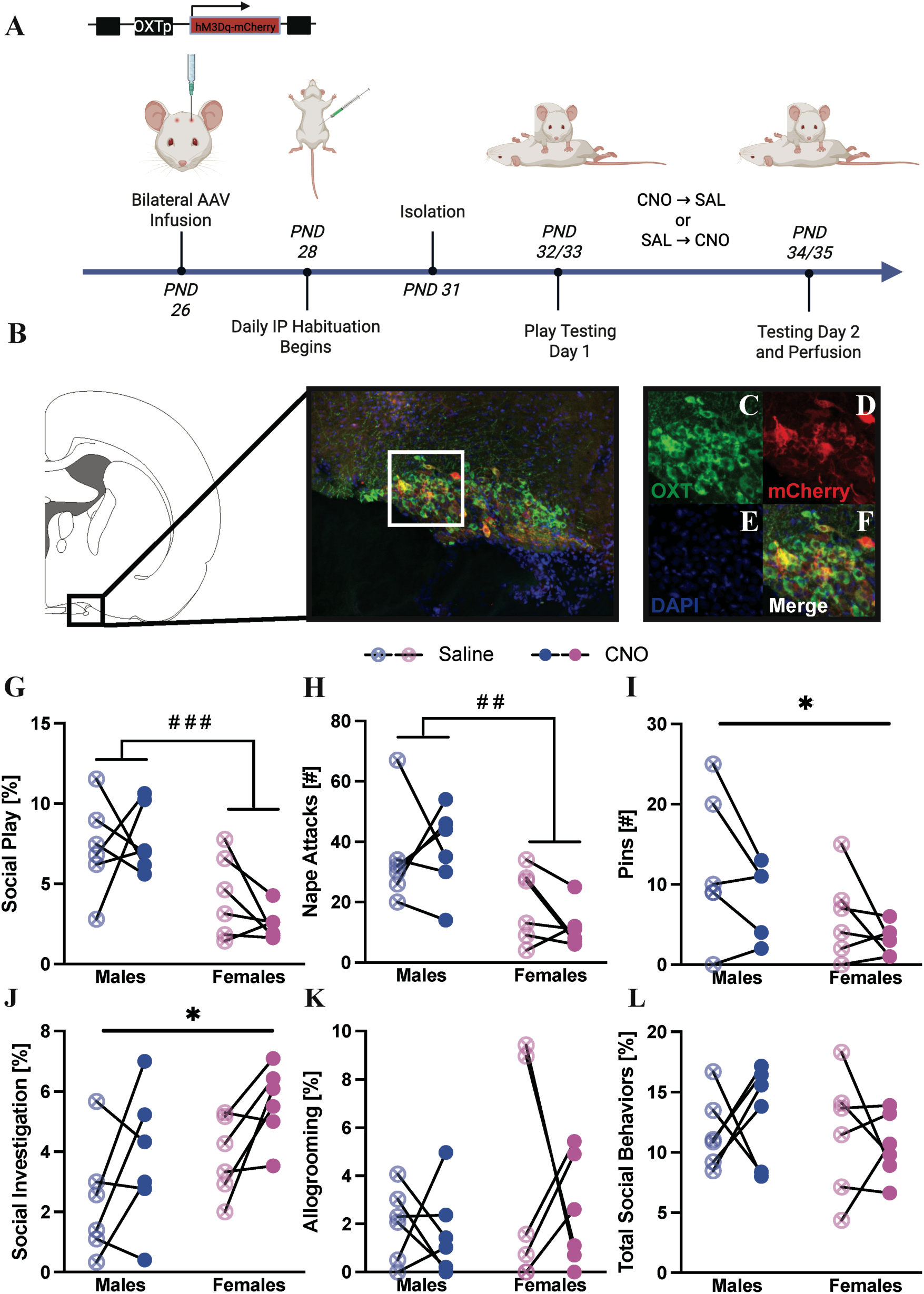
Chemogenetic stimulation of SON^OXT^ neurons decreases the number of pins and increases social investigation in juvenile male and female rats. (A) Timeline of experimental procedures in which juvenile rats (26-day-old) underwent stereotaxic surgery for infusion of AAV-OXTp-hM3Dq-mCherry into the SON. Following recovery and habituation to IP injections, all rats underwent two days of social play testing in which either saline or CNO was administered 30 min prior to testing. Drug conditions were randomly assigned and counterbalanced over the two days of testing. (B) Rat brain atlas image (modified from Paxinos and Watson, 2007) indicating the location of the SON and representative image of AAV-OXTp-hM3Dq-mCherry expression in OXT neurons of the SON as well as magnification of image denoting OXT (C, green) and mCherry (D, red) positive neurons that are co-localized using DAPI (E, blue) as a cellular marker. Irrespective of drug treatment, females showed lower duration of social play (G) and fewer nape attacks than males (H). CNO administration decreased the number of pins (I) and increased the duration of social investigation (J) in both sexes compared to saline administration. Neither drug nor sex altered the duration of allogrooming (K) and total duration of social behaviors (L). Durations of social play (G), social investigation (J), allogrooming (K), and total social behavior (L) are expressed as a percentage of total time. 2-way ANOVA main effect of drug, *: p < 0.05; main effect of sex, ##: p < 0.01; ###: p < 0.001.

**Table 3.** Experiment 2: Effects of chemogenetic stimulation of SON^OXT^ neurons; Two-way ANOVA statistics for behaviors quantified in social play testing. Significant main effects and interactions are notated in **bold**.

|  | Drug Treatment | Sex | Drug x Sex |
| --- | --- | --- | --- |
| Social Play [%] | $F_{1,10} = 0.36$,<br>$p = 0.56$, $\eta^2_p = 0.04$ | **$F_{1,10} = 30.4$,**<br>**$p = 0.0003$, $\eta^2_p = 0.61$** | $F_{1,10} = 1.11$,<br>$p = 0.32$, $\eta^2_p = 0.10$ |
| Nape Attacks [#] | $F_{1,10} = 0.30$,<br>$p = 0.60$, $\eta^2_p = 0.03$ | **$F_{1,10} = 13.3$,**<br>**$p = 0.005$, $\eta^2_p = 0.64$** | $F_{1,10} = 1.06$,<br>$p = 0.33$, $\eta^2_p = 0.10$ |
| Pins [#] | **$F_{1,10} = 5.97$,**<br>**$p = 0.03$, $\eta^2_p = 0.37$** | $F_{1,10} = 3.28$,<br>$p = 0.10$, $\eta^2_p = 0.54$ | $F_{1,10} = 0.28$,<br>$p = 0.61$, $\eta^2_p = 0.03$ |
| Social Investigation [%] | **$F_{1,10} = 16.8$,**<br>**$p = 0.02$, $\eta^2_p = 0.41$** | $F_{1,10} = 4.31$,<br>$p = 0.06$, $\eta^2_p = 0.43$ | $F_{1,10} = 0.08$,<br>$p = 0.79$, $\eta^2_p = 0.008$ |
| Allogrooming [%] | $F_{1,10} = 0.25$,<br>$p = 0.63$, $\eta^2_p = 0.02$ | $F_{1,10} = 1.52$,<br>$p = 0.25$, $\eta^2_p = 0.07$ | $F_{1,10} = 0.06$,<br>$p = 0.81$, $\eta^2_p = 0.006$ |
| Total Social Behavior [%] | $F_{1,10} = 0.04$,<br>$p = 0.85$, $\eta^2_p = 0.003$ | $F_{1,10} = 3.23$,<br>$p = 0.10$, $\eta^2_p = 0.06$ | $F_{1,10} = 0.52$,<br>$p = 0.49$, $\eta^2_p = 0.05$ |

### Experiment 3: PVN^OXT^ neurons predominantly innervate the anterior subregion of the NAc in juvenile male and female rats

There was a main effect of NAc subregion on PVN^OXT^ labeled fibers (Figure 3; F_4,28_ = 5.87, *p*= 0.0015). Post-hoc Holm-Sidak’s test for multiple comparisons revealed that the anterior region of the NAc core/shell showed a higher density of PVN^OXT^ labeled fibers compared to the intermediate NAc core (*p =* 0.012), intermediate NAc shell (*p =* 0.0021), and posterior NAc shell (*p=*0.0067). There were no main effects of sex (F_1,8_ = 0.23, *p*= 0.65) or interaction of sex x NAc subregion (F_4,28_ = 0.93, *p* = 0.46) on fiber density measures. Based on these results, the anterior subregion of the NAc was chosen as the region of interest for experiments 4-6.

**Figure 3.**
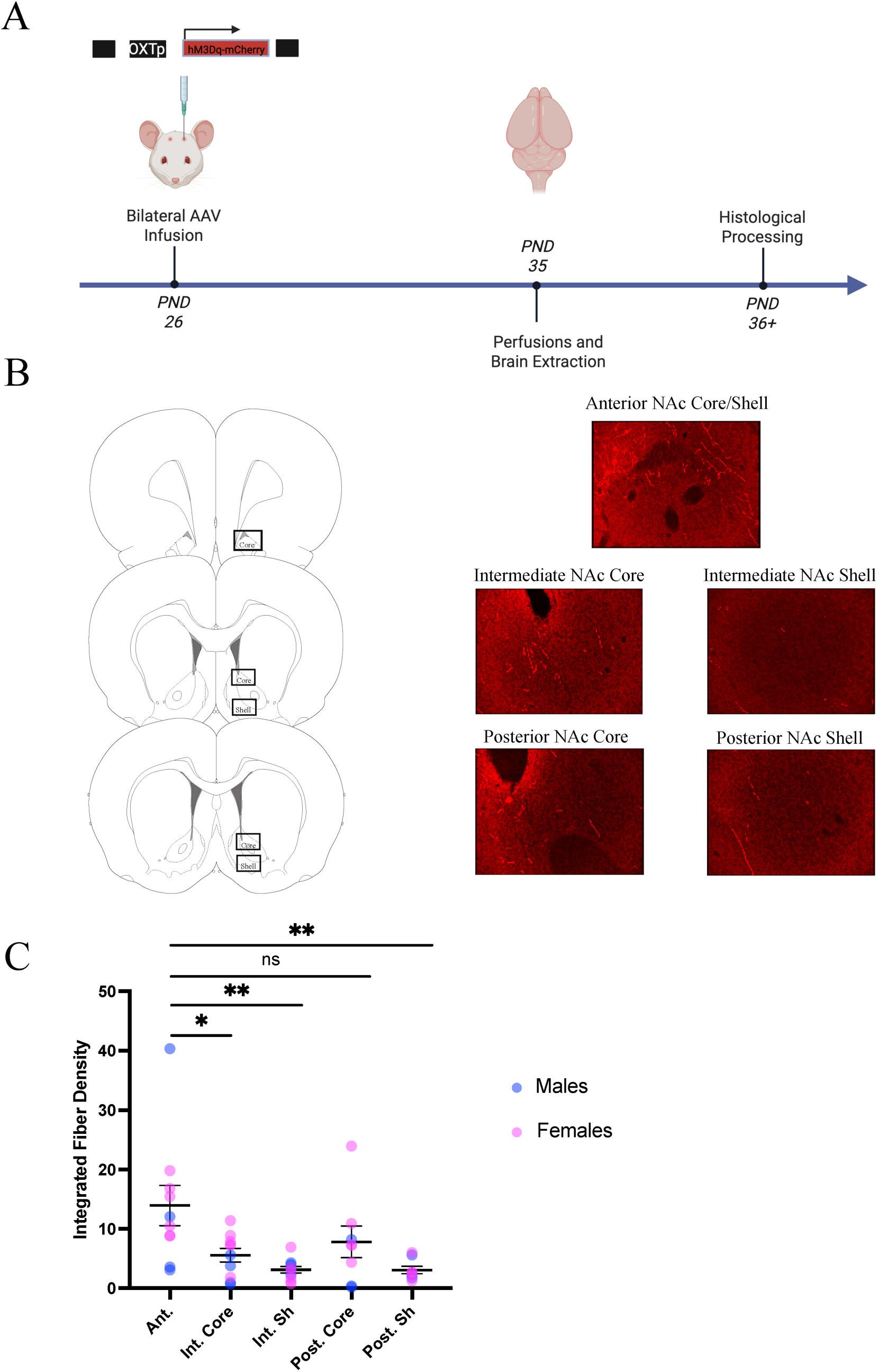
The anterior subregion of the NAc shows a higher integrated density of PVN^OXT^ fibers compared to the intermediate and posterior subregions of the NAc. (A) Schematic of experimental timeline in which juvenile male and female rats (22-days-old) were infused either AAV-OXTp-mCherry or AAV-OXTp-Venus into the PVN. At 35 days of age, rats were perfused and brains were collected for histological processing and OXT-immunoreactive fiber density analysis. (B) Imaging parameters and representative images for each of the five levels of the NAc that were analyzed. (C) The anterior subregion of the NAc (Ant.) shows a higher density of OXT-immunoreactive fibers originating from the PVN in comparison to the intermediate core (Int. Core), intermediate shell (Int. Sh), and the posterior shell (Post. Sh). 2-way ANOVA; main effect of NAc subregion, *: *p* < 0.05, **: *p* < 0.01, Holm’s-Sidak post-hoc tests.

### Experiment 4: Chemogenetic stimulation of PVN^OXT^ terminals in the NAc does not alter social play behaviors in juvenile male and female rats but increases social investigation in males

There were no effects of drug condition or sex on the duration of social play or number of nape attacks (See table 4 for statistical details). There was a main effect of sex on the number of pins, in which females displayed fewer pins than males, regardless of drug condition (Figure 4F). Additionally, there was a significant sex x drug interaction on social investigation (Figure 4G). Holm-Sidak post-hoc tests revealed that CNO administration significantly increased social investigation in males (*p* = 0.03) but tended to decrease social investigation in females (*p* = 0.06). There were no effects of sex or drug condition on the duration of allogrooming or cumulative duration of social behaviors (*p* > 0.05 for both; Figure 4). Additionally, local infusion of CNO outside of the NAc in PVN^OXT^ hM3Dq transfected rats did not alter any social or non-social behaviors in either sex (*p* > 0.05 for all, Supplemental Figure 3 B-G; see Supplemental Table 2 for statistical details).

**Figure 4.**
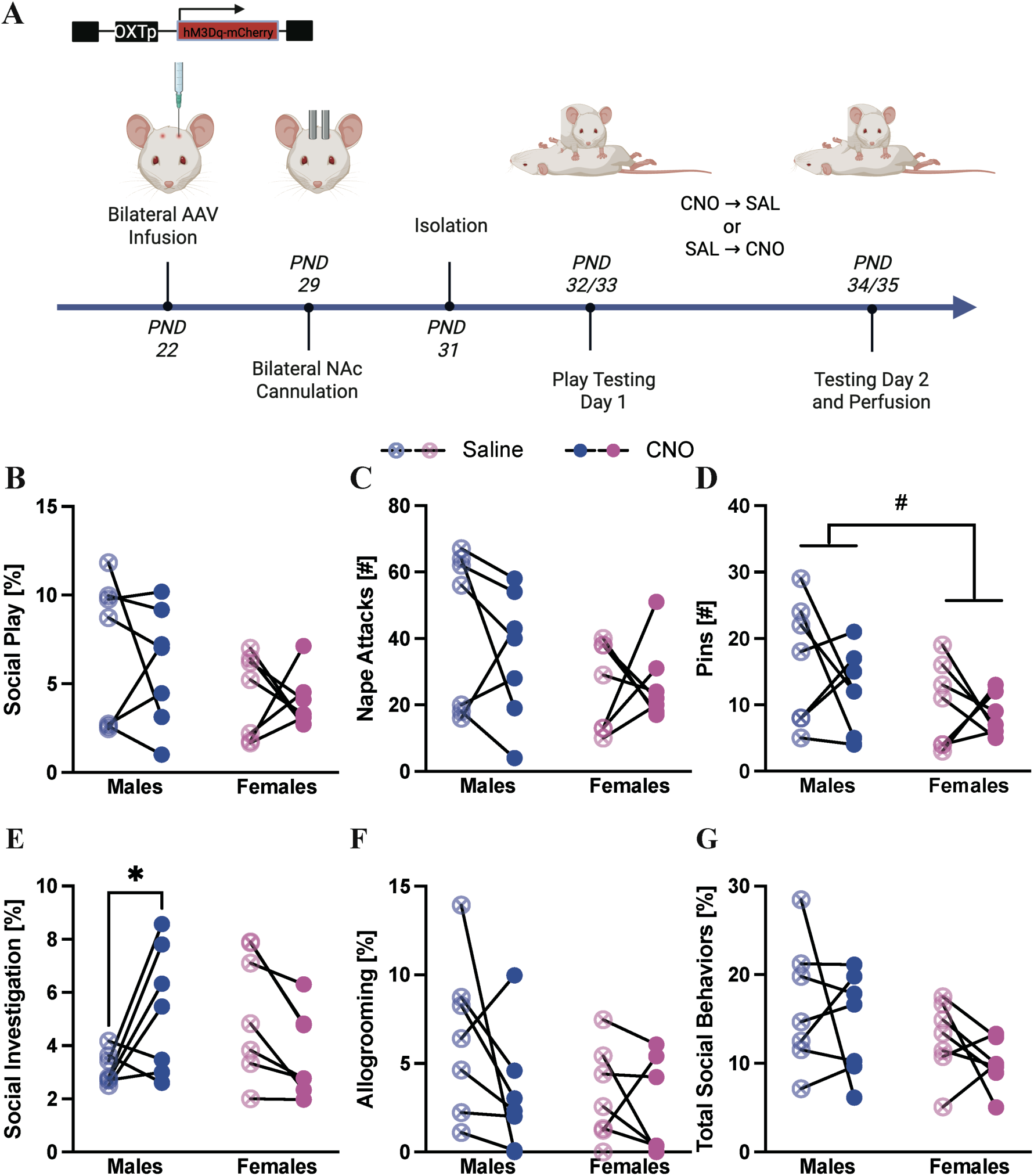
Chemogenetic stimulation of PVN^OXT^ terminals in the NAc does not alter social play behavior, but sex-specifically alters social investigation in juvenile rats. (A) Timeline of experimental procedures in which juvenile rats underwent stereotaxic surgery at 22 days of age for infusion of AAV-OXTp-hM3Dq-mCherry into the PVN and at 29 days of age for bilateral cannulation of the NAc. Following recovery and habituation to cannulae infusion procedures, rats underwent two days of behavioral testing in which either saline or CNO was locally infused in a counterbalanced manner 20 minutes prior to testing. (B-G) CNO administration did not alter the duration of social play (B), number of nape attacks (C), number of pins (D), duration of allogrooming (F), or cumulative duration of social behaviors (G). However, administration of CNO sex-specifically altered social investigation, wherein males showed a significant increase, and females a strong tendency towards a decrease, in duration of social investigation (E). Additionally, females showed fewer pins than males regardless of drug administration (D). Durations of social play (G), investigation (J), allogrooming (K), and combined social behaviors (L) are expressed as a percentage of total time. 2-way ANOVA; main effect of sex, #: *p* < 0.05; Sex x Drug interaction *: *p* < 0.05, Holm’s-Sidak post-hoc test.

**Table 4.** Experiment 4: Effects of chemogenetic stimulation of PVN^OXT^ terminals in the NAc; Two-way ANOVA statistics for behaviors quantified in social play testing. Significant main effects and interactions are notated in **bold**.

|  | Drug Treatment | Sex | Drug x Sex |
| --- | --- | --- | --- |
| Social Play [%] | $F_{1,12} = 0.37$,<br>$p = 0.56$, $\eta^2_p = 0.03$ | $F_{1,12} = 3.52$,<br>$p = 0.09$, $\eta^2_p = 0.29$ | $F_{1,12} = 0.04$,<br>$p = 0.84$, $\eta^2_p = 0.004$ |
| Nape Attacks [#] | $F_{1,12} = 0.43$,<br>$p = 0.52$, $\eta^2_p = 0.03$ | $F_{1,12} = 3.19$,<br>$p = 0.10$, $\eta^2_p = 0.29$ | $F_{1,12} = 0.50$,<br>$p = 0.50$, $\eta^2_p = 0.04$ |
| Pins [#] | $F_{1,12} = 1.12$,<br>$p = 0.31$, $\eta^2_p = 0.09$ | **$F_{1,12} = 5.02$,**<br>**$p = 0.04$, $\eta^2_p = 0.23$** | $F_{1,12} = 0.18$,<br>$p = 0.68$, $\eta^2_p = 0.02$ |
| Social Investigation [%] | $F_{1,12} = 0.29$,<br>$p = 0.60$, $\eta^2_p = 0.02$ | $F_{1,12} = 0.08$,<br>$p = 0.78$, $\eta^2_p = 0.02$ | **$F_{1,12} = 11.7$,**<br>**$p = 0.005$, $\eta^2_p = 0.49$** |
| Allogrooming [%] | $F_{1,12} = 2.14$,<br>$p = 0.17$, $\eta^2_p = 0.21$ | $F_{1,12} = 0.32$,<br>$p = 0.58$, $\eta^2_p = 0.20$ | $F_{1,12} = 1.19$,<br>$p = 0.30$, $\eta^2_p = 0.08$ |
| Total Social Behavior [%] | $F_{1,12} = 1.37$,<br>$p = 0.26$, $\eta^2_p = 0.10$ | $F_{1,12} = 4.66$,<br>$p = 0.052$, $\eta^2_p = 0.25$ | $F_{1,12} = 0.05$,<br>$p = 0.84$, $\eta^2_p = 0.004$ |

### Experiment 5: Acute OXT infusion into the NAc decreases social play behaviors in juvenile male rats and has no behavioral effects in juvenile female rats

There was a significant sex x drug interaction on the duration of social play, number of nape attacks, and number of pins (Figure 5B-D; see Table 5 for statistical details). Holm-Sidak post-hoc tests showed that infusion of the high dose of OXT into the NAc in males significantly decreased the duration of social play (5 µM: *p*=0.0018; Figure 5B), the number of nape attacks (5µM: *p*=0.01; Figure 5C) and number of pins (5µM: *p*=0.0025; Figure 5D) compared to saline infusion. The male specific effects of OXT administration eliminated the baseline sex difference in duration of social play and number of nape attacks, in which saline-treated females played less and showed fewer nape attacks than saline-treated males (*p* = 0.004 and *p* = 0.02, respectively) while OXT-treated males and females showed similar levels of social play (1µM: p = 0.09; 5µM: p = 0.21) and number of nape attacks (1µM: p = 0.25; 5µM: p = 0.25). No effect of sex or drug was found on the duration of social investigation, allogrooming, or cumulative duration of social behavior (*p* > 0.05 for all, Figure 5E-G).

**Figure 5.**
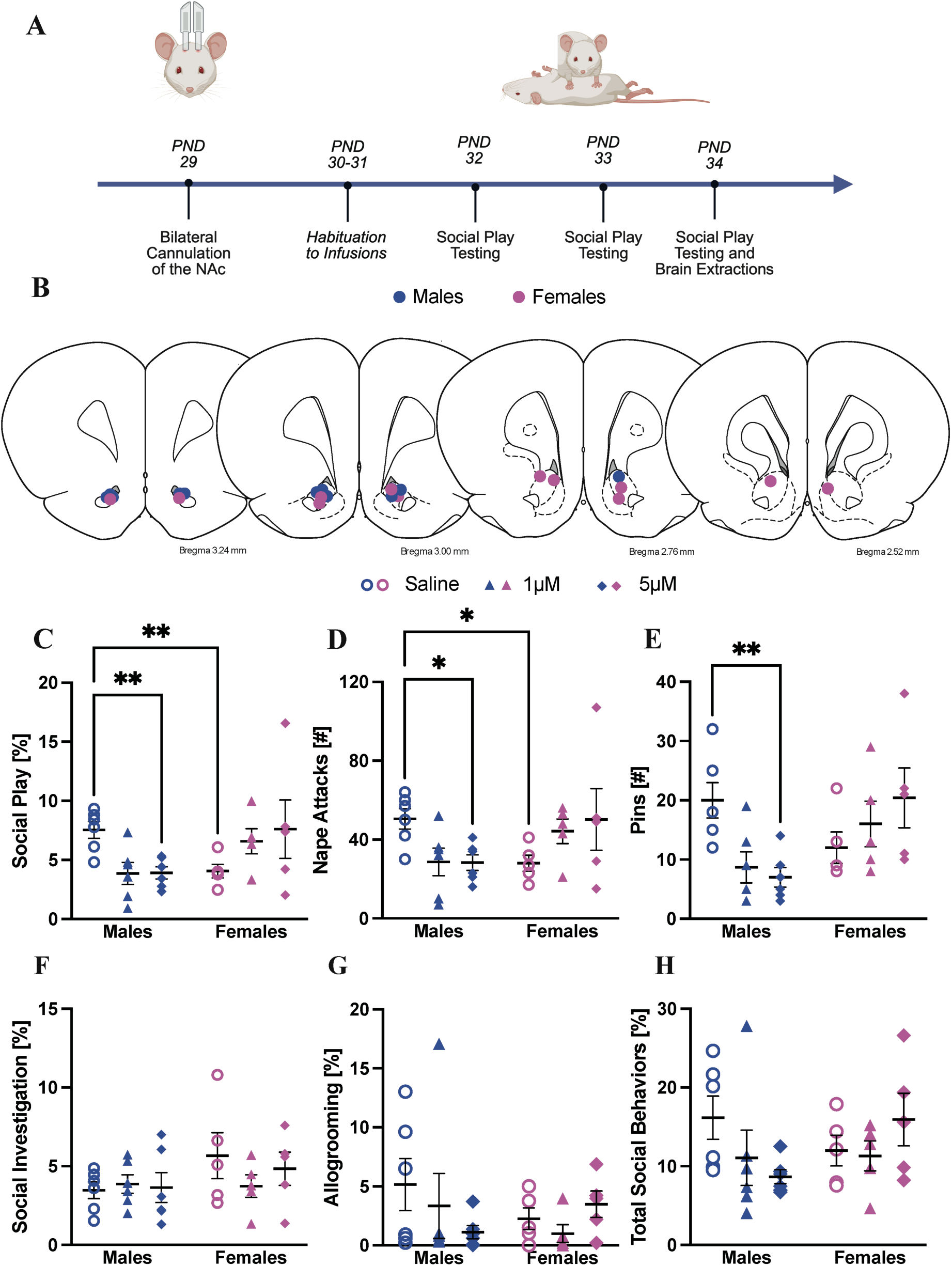
Acute infusion of OXT into the NAc sex-specifically alters social play behaviors in juvenile rats. (A) Timeline of experimental procedures in which juvenile rats underwent stereotaxic surgery at 28 days of age for bilateral cannulation of the NAc. Following recovery and habituation to cannulae infusion procedures, rats underwent three days of social play testing in which either saline, 1 µM OXT, or 5 µM OXT was locally infused om a counterbalanced manner 20 minutes prior to testing. (B) Schematic depicting the location of cannula placement within the NAc adapted from Paxinos and Watson (2007) (C-D) There was a baseline sex difference in which saline-treated females showed a shorter duration of social play and fewer nape attacks than saline-treated males. This sex difference was eliminated following OXT infusion, with males showing a decrease in duration of social play and number of nape attacks at the 5 µM dose of OXT. (E) Males showed a decrease in the number of pins at the 5 µM dose of OXT. There were no effects of OXT on the duration of social investigation (F), duration of allogrooming (G), or cumulative duration of social behavior (H). 2-way repeated measures ANOVA, Holm-Sidak post-hoc *: p < 0.05, **: p < 0.01.

**Table 5.** Experiment 5: Effects of OXT infusion in the NAc; Two-way ANOVA statistics for behaviors quantified in social play testing. Significant main effects and interactions are notated in **bold**.

|  | Drug Treatment | Sex | Drug x Sex |
| --- | --- | --- | --- |
| Social Play [%] | $F_{1.7, 15.5} = 0.15$,<br>$p = 0.84$, $\eta^2_p = 0.02$ | $F_{1,9} = 1.14$,<br>$p = 0.31$, $\eta^2_p = 0.05$ | **$F_{2,18} = 5.32$,**<br>**$p = 0.02$, $\eta^2_p = 0.37$** |
| Nape Attacks [#] | $F_{1.8, 15.9} = 0.10$,<br>$p = 0.89$, $\eta^2_p = 0.06$ | $F_{1,9} = 0.53$,<br>$p = 0.48$, $\eta^2_p = 0.22$ | **$F_{1.8, 15.9} = 5.28$,**<br>**$p = 0.02$, $\eta^2_p = 0.14$** |
| Pins [#] | $F_{1.8, 16.2} = 0.58$,<br>$p = 0.55$, $\eta^2_p = 0.06$ | $F_{1,9} = 3.92$,<br>$p = 0.08$, $\eta^2_p = 0.11$ | **$F_{2,18} = 5.16$,**<br>**$p = 0.02$, $\eta^2_p = 0.36$** |
| Social Investigation [%] | $F_{1.7, 15.4} = 0.49$,<br>$p = 0.59$, $\eta^2_p = 0.05$ | $F_{1,9} = 1.43$,<br>$p = 0.26$, $\eta^2_p = 0.14$ | $F_{2,18} = 1.12$,<br>$p = 0.35$, $\eta^2_p = 0.11$ |
| Allogrooming [%] | $F_{1.9, 16.9} = 0.65$,<br>$p = 0.52$, $\eta^2_p = 0.07$ | $F_{1,9} = 0.31$,<br>$p = 0.59$, $\eta^2_p = 0.03$ | $F_{2,18} = 1.89$,<br>$p = 0.18$, $\eta^2_p = 0.17$ |
| Total Social Behavior [%] | $F_{1.9, 17.5} = 0.77$,<br>$p = 0.47$, $\eta^2_p = 0.08$ | $F_{1,9} = 0.20$,<br>$p = 0.66$, $\eta^2_p = 0.02$ | $F_{2,18} = 3.01$,<br>$p = 0.07$, $\eta^2_p = 0.25$ |

### Experiment 6: Duration of social play correlates negatively with the proportion of *fos+* neurons and the proportion of activated *oxtr* mRNA-expressing neurons in the anterior NAc in both juvenile male and female rats

Females showed lower levels of social play behaviors, including lower duration of social play (t_(12)_ =4.14, *p =* 0.001, Figure 6A), fewer number of pins (t_(12)_ =2.60, *p =* 0.02, Figure 6C), and lower duration of total social behaviors (t_(12)_ =3.50, *p =* 0.004, Figure 6F) compared to males. There was no effect of sex on the number of nape attacks, duration of social investigation, and duration of allogrooming (*p* > 0.05 for all, Figure 6).

There were no main effects of sex, play condition, or interaction effects for *fos, oxtr,* or proportions of *fos+oxtr* mRNA in the anterior NAc (Figure 7 F-G; *p* > 0.05 for all; see Table 6 for statistical details).

There was a significant negative correlation between the duration of social play and the number of *fos*+ nuclei in the anterior NAc (r_(12)_ = -0.69, *p*= 0.007, Figure 7 F’) and between the duration of social play and the proportion of activated *oxtr*+ nuclei ([*oxtr+fos+/oxtr+*]*100; r_(12)_ = -0.67, *p*= 0.009, Figure 7 I’). When analyzed by sex, there was a significant negative correlation between the duration of social play and the proportion of activated *oxtr*+ nuclei in males (r_(6)_ = -0.73, *p*= 0.04, Figure 7 I’’) but not females (r_(4)_ = -0.71, *p* = 0.11, Figure 7 I’’’). No significant correlations were detected between duration of social play and *oxtr+fos+* nuclei or the proportion of activated nuclei co-expressing *oxtr* ([*oxtr+fos+/fos+*]*100; Figure 7 G-H, respectively). No significant correlations were detected between duration of social investigation and number of *fos+ nuclei*, *fos+oxtr*+ nuclei, the proportion of activated *oxtr+* nuclei, or the proportion of activated nuclei co-expressing *oxtr* (*p* > 0.05 for all, Supplemental Figure 4).

**Figure 6.**
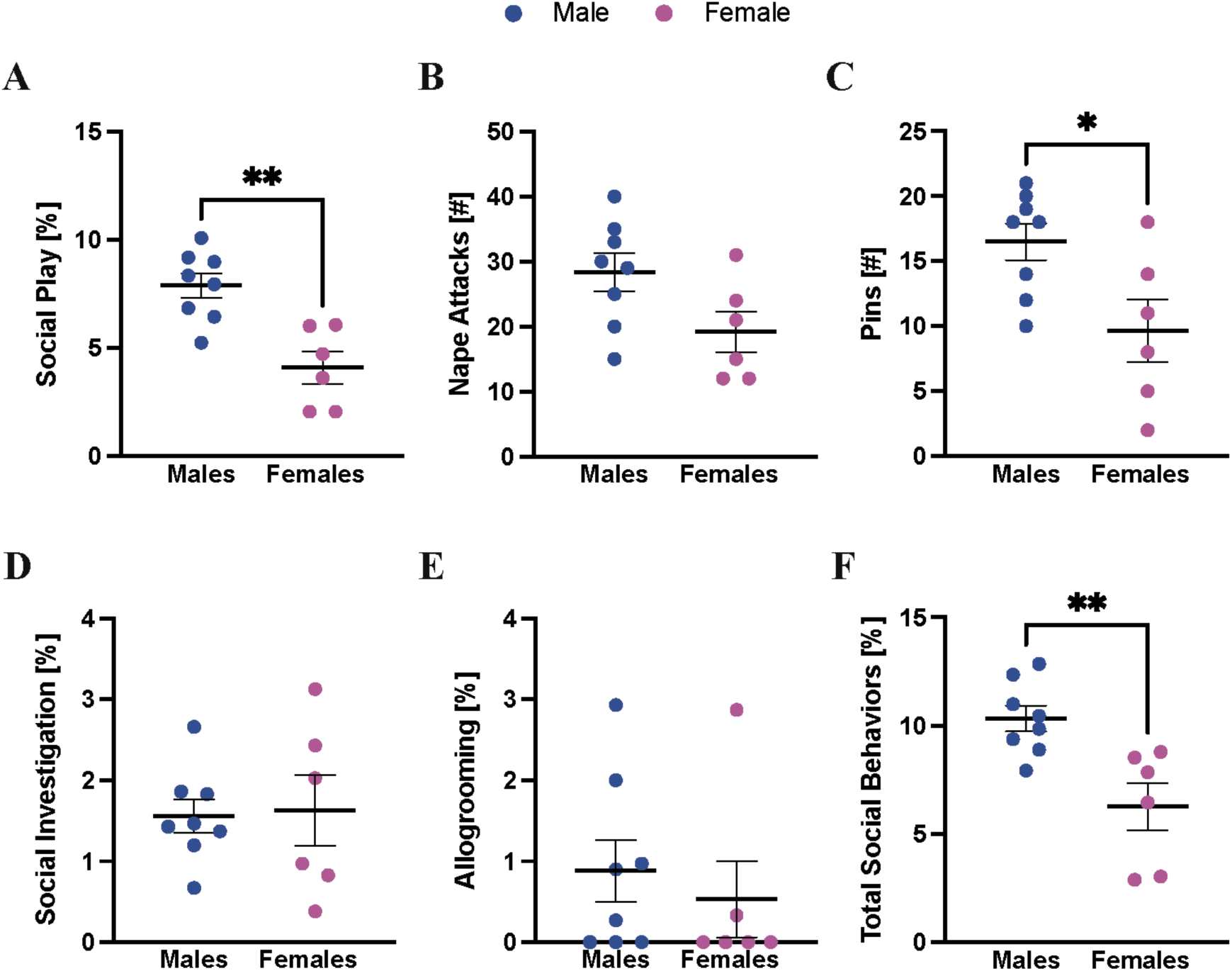
Juvenile female rats play at lower levels than juvenile male rats. Females played less (A), showed fewer pins (C), and had lower levels of total social behavior (F) than males during a 10-minute social play test. There were no differences between the sexes in the number of nape attacks (B), duration of social investigation (D) or duration of allogrooming (E). Durations of social play (A), investigation (D), allogrooming (E), and combined social behaviors (F) are expressed as a percentage of total time. Unpaired t-test; *: *p* < 0.05; **: *p* < 0.01.

**Figure 7.**
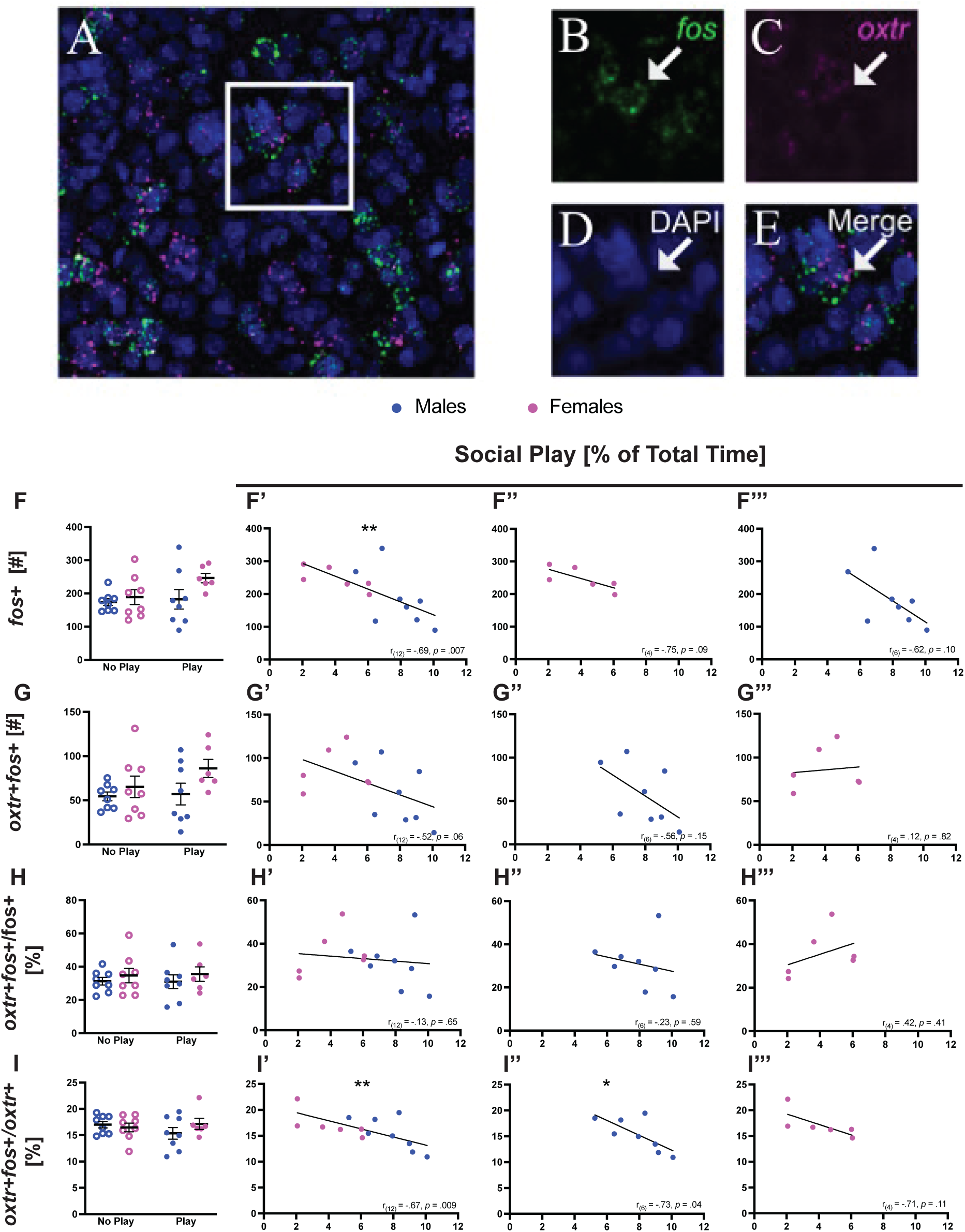
Exposure to social play does not alter activation patterns within the NAc in juvenile male and female rats but is negatively correlated with the number of activated *oxtr* mRNA+ neurons and the proportion of activated *oxtr* mRNA+ neurons in the anterior NAc (A) Representative image of *oxtr* and *fos* mRNA expressing nuclei in the NAc. (B-E) Examples of labeled nuclei from the white-boxed region of (A) positive for *fos* (B; green puncta), *oxtr* (C; magenta puncta), DAPI (D; blue nuclear stain), and double-labeled nuclei (E). Neither play condition nor sex altered the (F) average number of *fos* mRNA+ nuclei, (G) average number of *oxtr+fos* mRNA double-labeled nuclei, (H) the proportion of activated *oxtr* mRNA+ nuclei, or (I) the proportion of activated nuclei expressing *oxtr* mRNA. 2-way ANOVA; *p* > 0.05 for all. (F’) There is a significant negative correlation between duration of social play and the average number of *fos*+ nuclei, as well as (I’) duration of social play and the percentage of activated *oxtr+* nuclei. The association between social play and *fos+* nuclei was not detected in either males (F’’) or females (F’’’) alone, but there was a significant negative correlation between social play and the proportion of activated *oxtr+* nuclei in males only (I’’). No association between social play duration and average *oxtr+fos+* nuclei (G’-G’’’) or proportion of activated nuclei co-expressing *oxtr* (H’-H’’’) was detected.

**Table 6.** Experiment 6: Effects of social play exposure on activation patterns within the NAc; Two-way ANOVA statistics for *oxtr* and *fos* mRNA-expressing nuclei quantified. Significant main effects and interactions are notated in **bold**.

|  | Play Condition | Sex | Play x Sex |
| --- | --- | --- | --- |
| *fos* mRNA+ [#] | $F_{1,26} = 2.32$,<br>$p = 0.14$, $\eta^2_p = 0.08$ | $F_{1,26} = 3.36$,<br>$p = 0.08$, $\eta^2_p = 0.11$ | $F_{1,26} = 1.29$,<br>$p = 0.27$, $\eta^2_p = 0.05$ |
| *oxtr*+*fos* mRNA+ [#] | $F_{1,26} = 1.25$,<br>$p = 0.27$, $\eta^2_p = 0.05$ | $F_{1,26} = 3.58$,<br>$p = 0.07$, $\eta^2_p = 0.12$ | $F_{1,26} = 0.75$,<br>$p = 0.39$, $\eta^2_p = 0.03$ |
| *oxtr*+*fos*+/*oxtr*+ [%] | $F_{1,26} = 0.29$,<br>$p = 0.60$, $\eta^2_p = 0.01$ | $F_{1,26} = 0.46$,<br>$p = 0.50$, $\eta^2_p = 0.02$ | $F_{1,26} = 1.60$,<br>$p = 0.22$, $\eta^2_p = 0.06$ |
| *oxtr*+*fos*+/*fos*+ [%] | $F_{1,26} = 0.01$,<br>$p = 0.94$, $\eta^2_p = 0.0002$ | $F_{1,26} = 1.07$,<br>$p = 0.31$, $\eta^2_p = 0.04$ | $F_{1,26} = 0.02$,<br>$p = 0.88$, $\eta^2_p = 0.001$ |

## DISCUSSION

Here, we demonstrate the involvement of PVN^OXT^ neurons and OXT signaling within the NAc in the expression of social play in juvenile male rats. Chemogenetic stimulation of PVN^OXT^ neurons and intra-NAc infusions of OXT decreased the expression of social play behaviors in males, without altering the expression of social play behaviors in females. Additionally, the duration of social play, but not social investigation, correlated negatively with the proportion of activated *oxtr+* neurons in the NAc, an effect driven by males. This may provide first insights into the mechanism by which OXT in the NAc decreases social play in males. Lastly, we found that the duration of social investigation increased following chemogenetic stimulation of SON^OXT^ neurons (both sexes) and following chemogenetic stimulation of the PVN^OXT^ to NAc pathway (males only) without altering the most characteristic elements of social play behavior. This is the first study to suggest that activation of OXT neurons in the PVN as well as OXT signaling within the NAc sex-specifically modulate social play in juvenile rats, decreasing social play in males without altering social play in females.

Previous research showed that activation of PVN^OXT^ neurons, activation of PVN^OXT^ fibers into the NAc, and increased OXT signaling within the NAc all increased the expression of social behaviors in adult male or female rodent species (Dölen et al., 2013; He et al., 2021; Hou et al., 2023; Hung et al., 2017; Insel et al., 1998; Menon et al., 2018; Olazábal & Young, 2006b; H E Ross et al., 2009; Tang et al., 2020). Notably, most of these studies found increased social investigation or social interaction following such OXTergic manipulations (Dölen et al., 2013; Grund et al., 2019; Hou et al., 2023; Hung et al., 2017; Tang et al., 2020). In line with these data, we found that chemogenetic stimulation of SON^OXT^ neurons and of PVN^OXT^ terminals in the NAc increased social investigation in juvenile rats.

Yet, in contrast to our hypothesis, chemogenetic stimulation of PVN^OXT^ neurons as well as OXT infused into the NAc decreased the duration of social play and the number of nape attacks in juvenile male rats. This may be more in line with previous studies showing decreased expression of social behaviors following manipulations that increased OXT signaling (Beery, 2015; Duque-Wilckens et al., 2020; Higuchi et al., 2025; Shamay-Tsoory & Abu-Akel, 2016; Steinman et al., 2019). For example, chemogenetic stimulation of PVN^OXT^ neurons as well as chemogenetic stimulation of PVN^OXT^ neurons projecting to the bed nucleus of the stria terminalis (BNST) decreased social preference in adult male and female mice as measured in a three-chamber test (Higuchi et al., 2025). Moreover, OXT infusion into the BNST decreases social approach in adult male and female California mice (Duque-Wilckens et al., 2020). Thus, OXT may not indiscriminately increase the expression of social behaviors. These and other findings have led to the hypothesis that OXT enhances the salience of social information, thereby allowing the flexible expression of social behaviors (Shamay-Tsoory & Abu-Akel, 2016; Shamay-Tsoory et al., 2009; Steinman et al., 2019). Therefore, the direction of the effects of OXT on social behaviors may depend on internal and external contexts. In support, chemogenetic stimulation of PVN^OXT^ neurons increased social investigation towards a conspecific in adult female rats in a free but not in a caged interaction paradigm (Tang et al., 2020). Additionally, we previously showed that OXT infused in the lateral septum decreased social play duration in juvenile female rats when they were tested in their home cage but not when they were tested in a novel cage (Bredewold et al., 2014). It would therefore be of interest to determine whether our current findings showing OXT signaling decreasing social play behavior in juvenile male rats depends on the external context, such as testing environment or stimulus presentation.

A question that remains is how PVN^OXT^ neuronal signaling and OXT signaling in the NAc cause a decrease in social play behaviors in juvenile male rats. We found significant negative correlations between social play duration and NAc neuronal activation (as determined by fos) and between social play duration and the proportion of activated NAc^OXTR^ neurons in juvenile male rats. This suggests that an increase in the activation of NAc neurons, including NAc^OXTR^ neurons, contributed to the decrease in social play levels observed in juvenile male rats. If true, this may indicate that typical social play levels require suppressed activation of NAc output neurons. In support, pharmacological inhibition of the NAc increased social play in juvenile male rats (van Kerkhof et al., 2013). This concept is also in line with the prevailing view that inhibition of NAc neural output allows for the expression of rewarding behaviors (William A Carlezon & Thomas, 2009; Taha & Fields, 2006). For example, electrophysiological recordings demonstrate that a rewarding stimulus, intraoral administration of sucrose, inhibits NAc neurons in adult male rats (Roitman et al., 2005). Moreover, rats readily learn to self-administer a NMDA receptor antagonist into the NAc, which inhibits NAc neural output (W A Carlezon & Wise, 1996). Based on these collective data, we propose a model in which OXT activates NAc^OXTR^ neurons resulting in increased NAc output and a subsequent suppression of social play behaviors in juvenile male rats.

We found that chemogenetic stimulation of PVN^OXT^ neurons altered social play behaviors while chemogenetic stimulation of SON^OXT^ neurons altered social investigation. This may imply that these two hypothalamic OXT populations regulate different social behaviors. To the best of our knowledge, there are no other systematic studies dissecting the role of PVN^OXT^ and SON^OXT^ neuronal populations in the regulation of different behaviors. However, there is evidence suggesting that PVN^OXT^ and SON^OXT^ neuronal populations differentially modulating the same behaviors. For example, chemogenetic stimulation of PVN^OXT^ neurons decreased while chemogenetic stimulation of SON^OXT^ neurons increased food intake in adult male rats (Rea et al., 2025). Additionally, conditional knockout of the OXT gene prior to pregnancy in the SON, but not in the PVN, decreased the rate of pup survival in the offspring of female mice (Hagihara et al., 2023). These and our findings suggest the potential of different behavioral roles for OXT-producing neurons in the PVN versus the SON, which may, in part, be mediated through their distinct patterns of axonal projections within the brain (Hawthorn et al., 1985; Knobloch et al., 2012; Heather E Ross & Young, 2009).

Although chemogenetic stimulation of SON^OXT^ neurons decreased the number of pins and increased social investigation similarly in males and females, there were sex-specific effects of OXT manipulations in the other three experiments. Here, in contrast to males, chemogenetic stimulation of PVN^OXT^ neurons and intra-NAc infusions of OXT did not alter the expression of social play behaviors in females, nor did chemogenetic stimulation of the PVN^OXT^ to NAc pathway alter social investigation in females. These male-specific effects could be mediated by sex differences in OXT parameters in the PVN and NAc of juvenile rats. However, no sex differences were found in OXT mRNA expression in the PVN of adult rats nor in OXTR binding density in the NAc of juvenile rats (Dumais & Veenema, 2016; Dumais et al., 2013; Smith et al., 2017). Additionally, in the current study we found no sex difference in OXT fiber density in the NAc of juvenile rats. This extends previous work in juvenile rats showing no sex differences in OXT fiber density in six additional forebrain regions, including the anterior hypothalamus which contains dense OXT fibers that seem to form a tract that originates in the PVN. Although speculative, the absence of a sex difference in OXT fibers in the anterior hypothalamus that likely originate in the PVN may imply that there is no sex difference in OXT synthesis in the PVN of juvenile rats. Even in the absence of sex differences in these static OXT measures, there may be sex difference in baseline or social stimulus-induced OXT release. Although unknown in juvenile rats, microdialysis data in adult rats support this idea by revealing higher OXT release in the BNST in males versus females in response to exposure to a social stimulus (Dumais et al., 2016).

Another possible sex difference in the NAc-OXT system could involve OXTR coupling to different G protein subunits. Here, OXTR can couple and activate the inhibitory subunit G_i_ or the excitatory subunit G_q_ (Busnelli & Chini, 2018; Busnelli et al., 2012). OXTR-induced G_i_ versus G_q_ activation may produce different signaling effects, which, in turn, can have different effects on the expression of social behaviors. Whether this occurs in the NAc of rats is not known. But a first study in adult female California mice makes this plausible by showing that OXTR-G_i_ activation in the NAc modulates the expression of social approach but not social vigilance while OXTR-G_q_ activation modulates both social approach and social vigilance (Williams et al., 2020). It is further unknown whether there are sex differences in the coupling of OXTR to G_i_ or G_s_ in any species. However, such sex differences were found for other G protein-coupled receptors. For example, the corticotrophin releasing factor receptor can couple to distinct G proteins (Hillhouse & Grammatopoulos, 2006), with adult female rats showing higher G_q_ coupling than males in cortical tissue (Bangasser et al., 2010). A study in adult guinea pigs finds that sex differences in kappa-opioid receptor-G protein coupling are brain region specific (Wang et al., 2011). Several factors play a role in whether OXTR couples to G_i_ or G_q_, including the concentration of OXT (Busnelli & Chini, 2018). For example, *in vitro* studies showed that the OXTR-G_i_ complex is activated at higher OXT concentrations than the OXTR-G_q_ complex (Busnelli et al., 2012). Taken together, it would be interesting to determine whether sex differences exist in endogenous OXT release as well as in OXTR coupling to G_i_ or G_q_ subunits in the NAc. This knowledge may help reveal how increased OXT signaling in the NAc induced male-specific effects on social play behavior in juvenile rats.

Lastly, the lack of effects of the different OXT manipulations on social play behavior in juvenile female rats could be due to our gain-of-function approach for the DREADDs and pharmacological experiments. This gain-of-function approach is common in research on the role of OXT in human social behaviors as reflected by the numerous studies using intranasal OXT (reviewed in (Horta et al., 2020; Kraus et al., 2023; Mierop et al., 2020). These studies allow important insights into the potential of OXT to change the expression of social behaviors and may be informative for clinical studies (Quintana et al., 2021; Yin et al., 2024). Based on both human and rodent research, it has been argued that males and females may show different OXT dose-response relationships (Borland et al., 2019). Accordingly, using more than one dose of OXT in intranasal and intracerebral studies may be needed to uncover if behavioral effects of exogenous OXT are truly sex-specific or may depend on the dose (Procyshyn et al., 2024). Yet, our use of two doses of synthetic OXT infused into the NAc did not yield effects on social play behaviors in females. However, before excluding a role for the OXT system in the PVN and NAc in the regulation of social play behavior in females, future work may want to use a loss-of-function (inhibitory DREADDs, OXTR antagonists) approach. This will help uncover the necessity of endogenous OXT for the regulation of social play behavior, which may yield effects in females.

In conclusion, we demonstrated a role for PVN^OXT^ neurons and OXT signaling in the NAc in the regulation of social play behaviors in juvenile male but not female rats. The male-specific negative correlation between social play duration and the proportion of activated *oxtr+* neurons in the NAc, provides additional evidence for the sex-specific involvement of the NAc-OXT system in social play behavior. These findings led us to propose a model in males in which increased OXT signaling in the NAc facilitates the activation of the NAc, which in turn suppresses the expression of social play. Future research could test this further by determining whether inactivation of NAc^OXTR^ neurons increases the expression of social play in males and perhaps in females as well.

## References

Bangasser, D. A., Curtis, A., Reyes, B. A. S., Bethea, T. T., Parastatidis, I., Ischiropoulos, H., Van Bockstaele, E. J., & Valentino, R. J. (2010). Sex differences in corticotropin-releasing factor receptor signaling and trafficking: potential role in female vulnerability to stress-related psychopathology. Molecular Psychiatry, 15(9), 877, 896–904. 10.1038/mp.2010.66

Beery, A. K. (2015). Antisocial oxytocin: complex effects on social behavior. *Current Opinion in Behavioral Sciences*, 6, 174–182. 10.1016/j.cobeha.2015.11.006

Borland, J. M., Rilling, J. K., Frantz, K. J., & Albers, H. E. (2019). Sex-dependent regulation of social reward by oxytocin: an inverted U hypothesis. Neuropsychopharmacology, 44(1), 97–110. 10.1038/s41386-018-0129-2

Bredewold, R., Nascimento, N. F., Ro, G. S., Cieslewski, S. E., Reppucci, C. J., & Veenema, A. H. (2018). Involvement of dopamine, but not norepinephrine, in the sex-specific regulation of juvenile socially rewarding behavior by vasopressin. Neuropsychopharmacology, 43(10), 2109–2117. 10.1038/s41386-018-0100-2

Bredewold, R., Smith, C. J. W., Dumais, K. M., & Veenema, A. H. (2014). Sex-specific modulation of juvenile social play behavior by vasopressin and oxytocin depends on social context. Frontiers in Behavioral Neuroscience, 8, 216. 10.3389/fnbeh.2014.00216

Busnelli, M., & Chini, B. (2018). Molecular basis of oxytocin receptor signalling in the brain: what we know and what we need to know. Current Topics in Behavioral Neurosciences, 35, 3–29. 10.1007/7854_2017_6

Busnelli, M., Saulière, A., Manning, M., Bouvier, M., Galés, C., & Chini, B. (2012). Functional selective oxytocin-derived agonists discriminate between individual G protein family subtypes. The Journal of Biological Chemistry, 287(6), 3617–3629. 10.1074/jbc.M111.277178

Calcagnetti, D. J., & Schechter, M. D. (1992). Place conditioning reveals the rewarding aspect of social interaction in juvenile rats. Physiology & Behavior, 51(4), 667–672. 10.1016/0031-9384(92)90101-7

Carlezon, William A, & Thomas, M. J. (2009). Biological substrates of reward and aversion: a nucleus accumbens activity hypothesis. Neuropharmacology, *56 Suppl 1*(Suppl 1), 122–132. 10.1016/j.neuropharm.2008.06.075

Carlezon, W A, & Wise, R. A. (1996). Rewarding actions of phencyclidine and related drugs in nucleus accumbens shell and frontal cortex. The Journal of Neuroscience, *16*(9), 3112–3122. 10.1523/JNEUROSCI.16-09-03112.1996

Chevallier, C., Huguet, P., Happé, F., George, N., & Conty, L. (2013). Salient social cues are prioritized in autism spectrum disorders despite overall decrease in social attention. Journal of Autism and Developmental Disorders, 43(7), 1642–1651. 10.1007/s10803-012-1710-x

Chevallier, C., Kohls, G., Troiani, V., Brodkin, E. S., & Schultz, R. T. (2012). The social motivation theory of autism. Trends in Cognitive Sciences, *16*(4), 231–239. 10.1016/j.tics.2012.02.007

Dölen, G., Darvishzadeh, A., Huang, K. W., & Malenka, R. C. (2013). Social reward requires coordinated activity of nucleus accumbens oxytocin and serotonin. Nature, 501(7466), 179–184. 10.1038/nature12518

Dumais, K. M., Alonso, A. G., Immormino, M. A., Bredewold, R., & Veenema, A. H. (2016). Involvement of the oxytocin system in the bed nucleus of the stria terminalis in the sex-specific regulation of social recognition. Psychoneuroendocrinology, 64, 79–88. 10.1016/j.psyneuen.2015.11.007

Dumais, K. M., Bredewold, R., Mayer, T. E., & Veenema, A. H. (2013). Sex differences in oxytocin receptor binding in forebrain regions: correlations with social interest in brain region- and sex-specific ways. *Hormones and Behavior*, 64(4), 693–701. 10.1016/j.yhbeh.2013.08.012

Dumais, K. M., & Veenema, A. H. (2016). Vasopressin and oxytocin receptor systems in the brain: Sex differences and sex-specific regulation of social behavior. Frontiers in Neuroendocrinology, 40, 1–23. 10.1016/j.yfrne.2015.04.003

Duque-Wilckens, N., Torres, L. Y., Yokoyama, S., Minie, V. A., Tran, A. M., Petkova, S. P., Hao, R., Ramos-Maciel, S., Rios, R. A., Jackson, K., Flores-Ramirez, F. J., Garcia-Carachure, I., Pesavento, P. A., Iñiguez, S. D., Grinevich, V., & Trainor, B. C. (2020). Extrahypothalamic oxytocin neurons drive stress-induced social vigilance and avoidance. Proceedings of the National Academy of Sciences of the United States of America, 117(42), 26406–26413. 10.1073/pnas.2011890117

Floresco, S. B. (2015). The nucleus accumbens: an interface between cognition, emotion, and action. *Annual Review of Psychology*, CC, 25–52. 10.1146/annurev-psych-010213-115159

Grund, T., Tang, Y., Benusiglio, D., Althammer, F., Probst, S., Oppenländer, L., Neumann, I. D., & Grinevich, V. (2019). Chemogenetic activation of oxytocin neurons: Temporal dynamics, hormonal release, and behavioral consequences. Psychoneuroendocrinology, *106*, 77–84. 10.1016/j.psyneuen.2019.03.019

Hagihara, M., Miyamichi, K., & Inada, K. (2023). The importance of oxytocin neurons in the supraoptic nucleus for breastfeeding in mice. Plos One, 18(3), e0283152. 10.1371/journal.pone.0283152

Hawthorn, J., Ang, V. T., & Jenkins, J. S. (1985). Effects of lesions in the hypothalamic paraventricular, supraoptic and suprachiasmatic nuclei on vasopressin and oxytocin in rat brain and spinal cord. Brain Research, *346*(1), 51–57. 10.1016/0006-8993(85)91093-5

He, Z., Zhang, L., Hou, W., Zhang, X., Young, L. J., Li, L., Liu, L., Ma, H., Xun, Y., Lv, Z., Li, Y., Jia, R., Li, J., & Tai, F. (2021). Paraventricular nucleus oxytocin subsystems promote active paternal behaviors in mandarin voles. The Journal of Neuroscience, 41(31), 6699– 6713. 10.1523/JNEUROSCI.2864-20.2021

Higuchi, Y., Ozawa, A., Kobayashi, R., Konno, T., & Arakawa, H. (2025). Functional disruption of oxytocin projections participates atypical social and anxiety-like behaviours in BTBR mouse model of autism. Open Biology, 15(8), 240387. 10.1098/rsob.240387

Hillhouse, E. W., & Grammatopoulos, D. K. (2006). The molecular mechanisms underlying the regulation of the biological activity of corticotropin-releasing hormone receptors: implications for physiology and pathophysiology. Endocrine Reviews, 27(3), 260–286. 10.1210/er.2005-0034

Horta, M., Pehlivanoglu, D., & Ebner, N. C. (2020). The role of intranasal oxytocin on social cognition: an integrative human lifespan approach. Current Behavioral Neuroscience Reports, 7(4), 175–192. 10.1007/s40473-020-00214-5

Hou, W., Huang, S., Li, L., Guo, X., He, Z., Shang, S., Jia, Z., Zhang, L., Qu, Y., Huang, C., Li, Y., Li, Y., Lv, Z., & Tai, F. (2023). Oxytocin treatments or activation of the paraventricular nucleus-the shell of nucleus accumbens pathway reduce adverse effects of chronic social defeat stress on emotional and social behaviors in Mandarin voles. Neuropharmacology, 230, 109482. 10.1016/j.neuropharm.2023.109482

Hung, L. W., Neuner, S., Polepalli, J. S., Beier, K. T., Wright, M., Walsh, J. J., Lewis, E. M., Luo, L., Deisseroth, K., Dölen, G., & Malenka, R. C. (2017). Gating of social reward by oxytocin in the ventral tegmental area. Science, 357(6358), 1406–1411. 10.1126/science.aan4994

Insel, T. R., & Shapiro, L. E. (1992). Oxytocin receptor distribution reflects social organization in monogamous and polygamous voles. Proceedings of the National Academy of Sciences of the United States of America, *89*(13), 5981–5985. 10.1073/pnas.89.13.5981

Insel, T. R., Winslow, J. T., Wang, Z., & Young, L. J. (1998). Oxytocin, vasopressin, and the neuroendocrine basis of pair bond formation. Advances in Experimental Medicine and Biology, *449*, 215–224. 10.1007/978-1-4615-4871-3_28

Jordan, R. (2003). Social play and autistic spectrum disorders: a perspective on theory, implications and educational approaches. Autism: The International Journal of Research and Practice, 7(4), 347–360. 10.1177/1362361303007004002

Keebaugh, A. C., & Young, L. J. (2011). Increasing oxytocin receptor expression in the nucleus accumbens of pre-pubertal female prairie voles enhances alloparental responsiveness and partner preference formation as adults. *Hormones and Behavior*, 60(5), 498–504. 10.1016/j.yhbeh.2011.07.018

Klawonn, A. M., & Malenka, R. C. (2018). Nucleus accumbens modulation in reward and aversion. Cold Spring Harbor Symposia on Quantitative Biology, 83, 119–129. 10.1101/sqb.2018.83.037457

Knobloch, H. S., Charlet, A., Hoffmann, L. C., Eliava, M., Khrulev, S., Cetin, A. H., Osten, P., Schwarz, M. K., Seeburg, P. H., Stoop, R., & Grinevich, V. (2012). Evoked axonal oxytocin release in the central amygdala attenuates fear response. Neuron, 73(3), 553–566. 10.1016/j.neuron.2011.11.030

Kraus, J., Výborová, E., & Silani, G. (2023). The effect of intranasal oxytocin on social reward processing in humans: a systematic review. Frontiers in Psychiatry, 14, 1244027. 10.3389/fpsyt.2023.1244027

Liu, Y., & Wang, Z. X. (2003). Nucleus accumbens oxytocin and dopamine interact to regulate pair bond formation in female prairie voles. Neuroscience, 121(3), 537–544. 10.1016/s0306-4522(03)00555-4

Menon, R., Grund, T., Zoicas, I., Althammer, F., Fiedler, D., Biermeier, V., Bosch, O. J., Hiraoka, Y., Nishimori, K., Eliava, M., Grinevich, V., & Neumann, I. D. (2018). Oxytocin Signaling in the Lateral Septum Prevents Social Fear during Lactation. Current Biology, 28(7), 1066–1078.e6. 10.1016/j.cub.2018.02.044

Mierop, A., Mikolajczak, M., Stahl, C., Béna, J., Luminet, O., Lane, A., & Corneille, O. (2020). How can intranasal oxytocin research be trusted? A systematic review of the interactive effects of intranasal oxytocin on psychosocial outcomes. Perspectives on Psychological Science, 15(5), 1228–1242. 10.1177/1745691620921525

Mitre, M., Marlin, B. J., Schiavo, J. K., Morina, E., Norden, S. E., Hackett, T. A., Aoki, C. J., Chao, M. V., & Froemke, R. C. (2016). A distributed network for social cognition enriched for oxytocin receptors. The Journal of Neuroscience, 36(8), 2517–2535. 10.1523/JNEUROSCI.2409-15.2016

Oettl, L.-L., Ravi, N., Schneider, M., Scheller, M. F., Schneider, P., Mitre, M., da Silva Gouveia, M., Froemke, R. C., Chao, M. V., Young, W. S., Meyer-Lindenberg, A., Grinevich, V., Shusterman, R., & Kelsch, W. (2016). Oxytocin enhances social recognition by modulating cortical control of early olfactory processing. *Neuron*, 90(3), 609–621. 10.1016/j.neuron.2016.03.033

Olazábal, D. E., & Young, L. J. (2006a). Species and individual differences in juvenile female alloparental care are associated with oxytocin receptor density in the striatum and the lateral septum. Hormones and Behavior, *49*(5), 681–687. 10.1016/j.yhbeh.2005.12.010

Olazábal, D. E., & Young, L. J. (2006b). Oxytocin receptors in the nucleus accumbens facilitate “spontaneous” maternal behavior in adult female prairie voles. Neuroscience, 141(2), 559–568. 10.1016/j.neuroscience.2006.04.017

Panksepp, J. (1981). The ontogeny of play in rats. Developmental Psychobiology, 14(4), 327–332. 10.1002/dev.420140405

Paxinos, G., & Watson, C. (2013). *The Rat Brain in Stereotaxic Coordinates* (7th Edition, p. 472). Academic Press. 10.1016/C2009-0-63235-9

Pellegrini, Anthony D., & Smith, P. K. (1998). The development of play during childhood: forms and possible functions. Child Psychology and Psychiatry Review, 3(2), 51–57. 10.1017/S1360641798001476

Pellegrini, Anthony D., & Smith, P. K. (Eds.). (2005). The Nature of Play: Great Apes and Humans (illustrated ed.). Guilford Press.

Pellegrini, A D, & Smith, P. K. (1998). Physical activity play: the nature and function of a neglected aspect of playing. *Child Development*, CS(3), 577–598. 10.1111/j.1467-8624.1998.tb06226.x

Pellegrini, A D. (1989). Elementary school children’s rough-and-tumble play. Early Childhood Research Quarterly, 4(2), 245–260. 10.1016/S0885-2006(89)80006-7

Pellis, S. M., & Pellis, V. C. (1997). The prejuvenile onset of play fighting in laboratory rats (Rattus norvegicus). Developmental Psychobiology, 31(3), 193–205. 10.1002/(SICI)1098-2302(199711)31:3<193::AID-DEV4>3.0.CO;2-N

Procyshyn, T. L., Dupertuys, J., & Bartz, J. A. (2024). Neuroimaging and behavioral evidence of sex-specific effects of oxytocin on human sociality. Trends in Cognitive Sciences. 10.1016/j.tics.2024.06.010

Quintana, D. S., Lischke, A., Grace, S., Scheele, D., Ma, Y., & Becker, B. (2021). Advances in the field of intranasal oxytocin research: lessons learned and future directions for clinical research. Molecular Psychiatry, 26(1), 80–91. 10.1038/s41380-020-00864-7

Rea, J. J., Liu, C. M., Hayes, A. M. R., Ohan, R., Schwartz, G. M., Bashaw, A. G., Klug, M. E., Decarie-Spain, L., Park, Y., Kao, A. E., Grinevich, V., & Kanoski, S. E. (2025). Oxytocin neurons in the paraventricular and supraoptic hypothalamic nuclei bidirectionally modulate food intake. Molecular Metabolism, 102220. 10.1016/j.molmet.2025.102220

Roitman, M. F., Wheeler, R. A., & Carelli, R. M. (2005). Nucleus accumbens neurons are innately tuned for rewarding and aversive taste stimuli, encode their predictors, and are linked to motor output. Neuron, 45(4), 587–597. 10.1016/j.neuron.2004.12.055

Ross, Heather E, Freeman, S. M., Spiegel, L. L., Ren, X., Terwilliger, E. F., & Young, L. J. (2009). Variation in oxytocin receptor density in the nucleus accumbens has differential effects on affiliative behaviors in monogamous and polygamous voles. The Journal of Neuroscience, 29(5), 1312–1318. 10.1523/JNEUROSCI.5039-08.2009

Ross, Heather E, & Young, L. J. (2009). Oxytocin and the neural mechanisms regulating social cognition and affiliative behavior. Frontiers in Neuroendocrinology, 30(4), 534–547. 10.1016/j.yfrne.2009.05.004

Ross, H E, Cole, C. D., Smith, Y., Neumann, I. D., Landgraf, R., Murphy, A. Z., & Young, L. J. (2009). Characterization of the oxytocin system regulating affiliative behavior in female prairie voles. Neuroscience, *162*(4), 892–903. 10.1016/j.neuroscience.2009.05.055

Salgado, S., & Kaplitt, M. G. (2015). The nucleus accumbens: A comprehensive review. *Stereotactic and Functional Neurosurgery*, 93(2), 75–93. 10.1159/000368279

Sawchenko, P. E., & Swanson, L. W. (1983). The organization of forebrain afferents to the paraventricular and supraoptic nuclei of the rat. The Journal of Comparative Neurology, 218(2), 121–144. 10.1002/cne.902180202

Shamay-Tsoory, S. G., & Abu-Akel, A. (2016). The social salience hypothesis of oxytocin. Biological Psychiatry, 79(3), 194–202. 10.1016/j.biopsych.2015.07.020

Shamay-Tsoory, S. G., Fischer, M., Dvash, J., Harari, H., Perach-Bloom, N., & Levkovitz, Y. (2009). Intranasal administration of oxytocin increases envy and schadenfreude (gloating). *Biological Psychiatry*, CC(9), 864–870. 10.1016/j.biopsych.2009.06.009

Shimada, M., & Sueur, C. (2018). Social play among juvenile wild Japanese macaques (Macaca fuscata) strengthens their social bonds. American Journal of Primatology, 80(1). 10.1002/ajp.22728

Smith, C. J. W., DiBenedictis, B. T., & Veenema, A. H. (2019). Comparing vasopressin and oxytocin fiber and receptor density patterns in the social behavior neural network: Implications for cross-system signaling. Frontiers in Neuroendocrinology, 53, 100737. 10.1016/j.yfrne.2019.02.001

Smith, C. J. W., Poehlmann, M. L., Li, S., Ratnaseelan, A. M., Bredewold, R., & Veenema, A. H. (2017). Age and sex differences in oxytocin and vasopressin V1a receptor binding densities in the rat brain: focus on the social decision-making network. Brain Structure & Function, 222(2), 981–1006. 10.1007/s00429-016-1260-7

Steinman, M. Q., Duque-Wilckens, N., & Trainor, B. C. (2019). Complementary neural circuits for divergent effects of oxytocin: social approach versus social anxiety. Biological Psychiatry, 85(10), 792–801. 10.1016/j.biopsych.2018.10.008

Taha, S. A., & Fields, H. L. (2006). Inhibitions of nucleus accumbens neurons encode a gating signal for reward-directed behavior. The Journal of Neuroscience, *26*(1), 217–222. 10.1523/JNEUROSCI.3227-05.2006

Tang, Y., Benusiglio, D., Lefevre, A., Hilfiger, L., Althammer, F., Bludau, A., Hagiwara, D., Baudon, A., Darbon, P., Schimmer, J., Kirchner, M. K., Roy, R. K., Wang, S., Eliava, M., Wagner, S., Oberhuber, M., Conzelmann, K. K., Schwarz, M., Stern, J. E., … Grinevich, V. (2020). Social touch promotes interfemale communication via activation of parvocellular oxytocin neurons. Nature Neuroscience, 23(9), 1125–1137. 10.1038/s41593-020-0674-y

Trezza, V., Baarendse, P. J. J., & Vanderschuren, L. J. M. J. (2010). The pleasures of play: pharmacological insights into social reward mechanisms. Trends in Pharmacological Sciences, 31(10), 463–469. 10.1016/j.tips.2010.06.008

Vanderschuren, L. J. M. J., Achterberg, E. J. M., & Trezza, V. (2016). The neurobiology of social play and its rewarding value in rats. Neuroscience and Biobehavioral Reviews, 70, 86–105. 10.1016/j.neubiorev.2016.07.025

van den Berg, C. L., Hol, T., Van Ree, J. M., Spruijt, B. M., Everts, H., & Koolhaas, J. M. (1999). Play is indispensable for an adequate development of coping with social challenges in the rat. Developmental Psychobiology, 34(2), 129–138.

van Kerkhof, L. W. M., Damsteegt, R., Trezza, V., Voorn, P., & Vanderschuren, L. J. M. J. (2013). Social play behavior in adolescent rats is mediated by functional activity in medial prefrontal cortex and striatum. Neuropsychopharmacology, 38(10), 1899–1909. 10.1038/npp.2013.83

Wang, Y.-J., Rasakham, K., Huang, P., Chudnovskaya, D., Cowan, A., & Liu-Chen, L.-Y. (2011). Sex difference in κ-opioid receptor (KOPR)-mediated behaviors, brain region KOPR level and KOPR-mediated guanosine 5’-O-(3-[35S]thiotriphosphate) binding in the guinea pig. The Journal of Pharmacology and Experimental Therapeutics, *339*(2), 438–450. 10.1124/jpet.111.183905

Williams, A. V., Duque-Wilckens, N., Ramos-Maciel, S., Campi, K. L., Bhela, S. K., Xu, C. K., Jackson, K., Chini, B., Pesavento, P. A., & Trainor, B. C. (2020). Social approach and social vigilance are differentially regulated by oxytocin receptors in the nucleus accumbens. Neuropsychopharmacology, 45(9), 1423–1430. 10.1038/s41386-020-0657-4

Yin, H., Jiang, M., Han, T., & Xu, X. (2024). Intranasal oxytocin as a treatment for anxiety and autism: From subclinical to clinical applications. Peptides, *176*, 171211. 10.1016/j.peptides.2024.171211

Young, L. J., Lim, M. M., Gingrich, B., & Insel, T. R. (2001). Cellular mechanisms of social attachment. Hormones and Behavior, 40(2), 133–138. 10.1006/hbeh.2001.1691

Yu, C. J., Zhang, S. W., & Tai, F. D. (2016). Effects of nucleus accumbens oxytocin and its antagonist on social approach behavior. Behavioural Pharmacology, 27(8), 672–680. 10.1097/FBP.0000000000000212

