## Supplemental Figures and Table for "Sex-specific regulation of social play in juvenile rats by oxytocin neurons in the hypothalamus and oxytocin signaling in the nucleus accumbens"

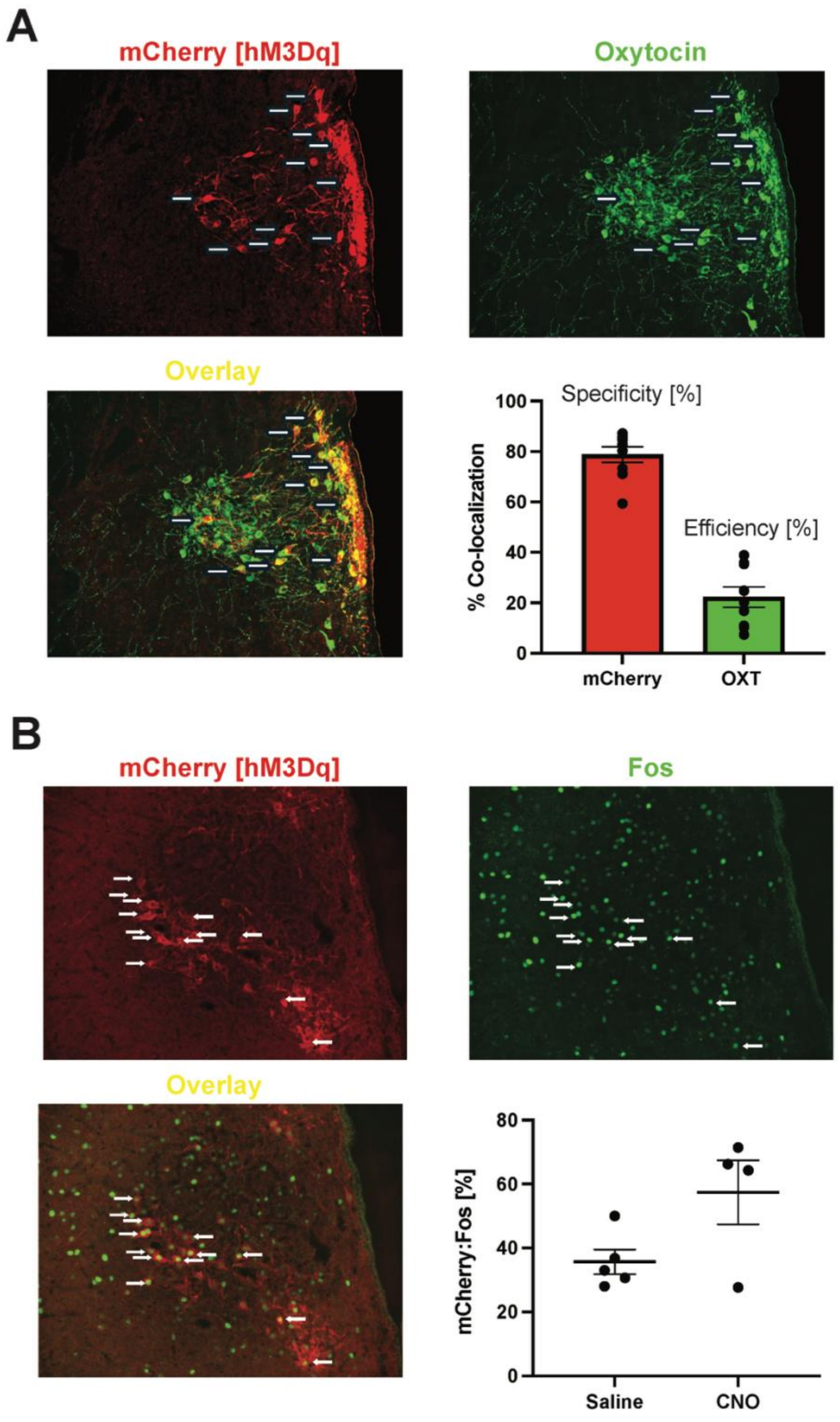

**Supplemental Figure 1. Validation of AAV1/2-OXTp-hM3Dq-mCherry in the PVN.** (A) Representative image of mCherry (red), OXT (green), and co-localization of mCherry and OXT (yellow) in the PVN used to determine the specificity (% of mCherry-ir cells co-localized in OXT-ir cells; 78.8%, SEM± 3.06) and efficiency (% of OXT-ir cells co-localized in mCherry-ir cells; 22.27%, SEM± 4.03). (B) Representative image of mCherry (red), Fos (green), and co-localization of mCherry and Fos (yellow) used to determine percentage of activated mCherry-ir+ neurons following administration of either saline (35.72%, SEM± 3.85) or CNO (57.43%, SEM± 10.02).

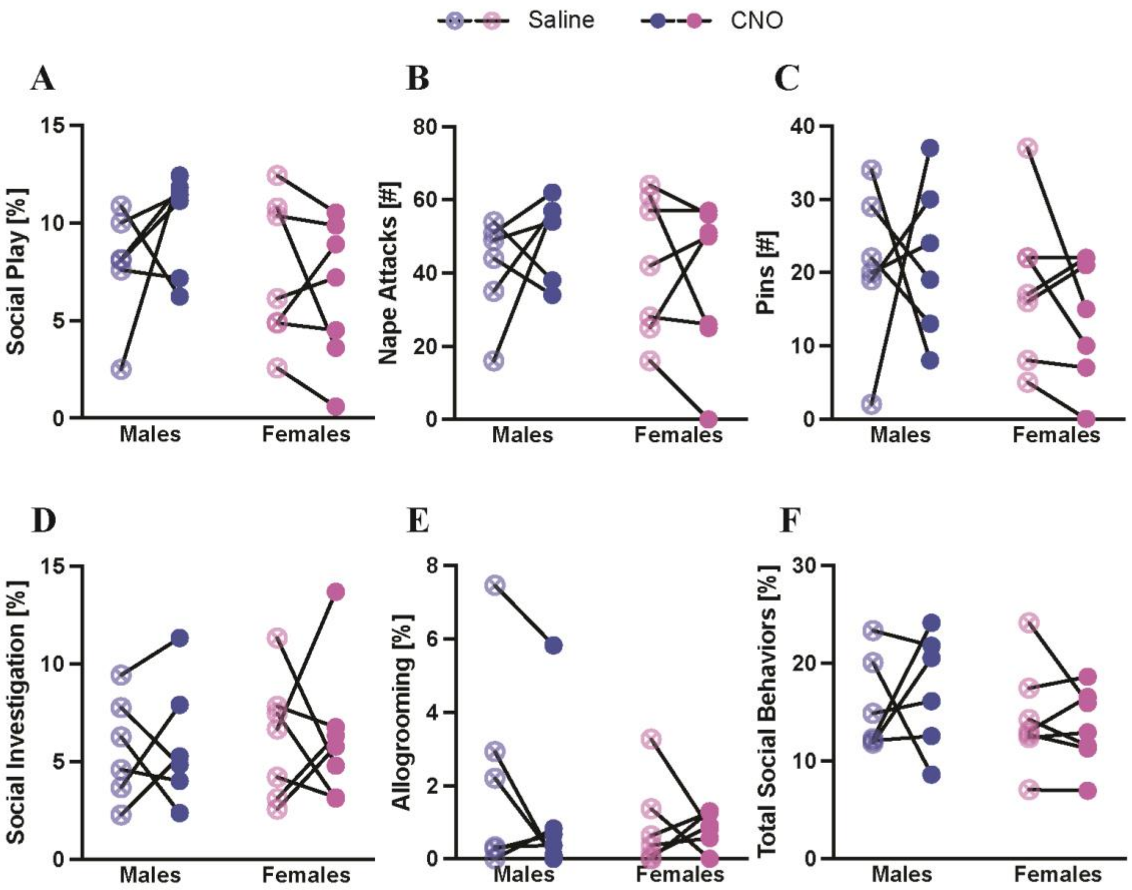


**Supplemental Figure 2. CNO administration in the absence of hM3Dq expression in the PVN does not alter social behaviors in juvenile male and female rats**. (A-F) CNO administration did not alter the duration of social play (A), number of nape attacks (B), number of pins (C), duration of social investigation (D), duration of allogrooming (E), or total duration of social behaviors (F) in juvenile male and female rats infused with AAV-OXTp-mCherry into the PVN. Social play, investigation, allogrooming and combined social behaviors are expressed as a percentage of total time.

**Supplemental Table 1.** Experiment 1: Effects of systemic CNO administration in the absence of hM3Dq; Two-way ANOVA statistics for behaviors quantified in social play testing. Significant main effects and interactions are notated in **bold**.

| | Drug Treatment | Sex | Drug x Sex |
|---|---|---|---|
| Social Play [%] | $F_{1,11} = 0.27$, $p = 0.61$, $\eta^2_p = 0.02$ | $F_{1,11} = 1.93$, $p = 0.19$, $\eta^2_p = 0.22$ | $F_{1,11} = 1.88$, $p = 0.20$, $\eta^2_p = 0.15$ |
| Nape Attacks [#] | $F_{1,11} = 0.19$, $p = 0.67$, $\eta^2_p = 0.02$ | $F_{1,11} = 0.59$, $p = 0.46$, $\eta^2_p = 0.10$ | $F_{1,11} = 1.32$, $p = 0.28$, $\eta^2_p = 0.11$ |
| Pins [#] | $F_{1,11} = 0.15$, $p = 0.70$, $\eta^2_p = 0.01$ | $F_{1,11} = 2.35$, $p = 0.15$, $\eta^2_p = 0.12$ | $F_{1,11} = 0.33$, $p = 0.58$, $\eta^2_p = 0.03$ |
| Social Investigation [%] | $F_{1,11} = 0.02$, $p = 0.88$, $\eta^2_p = 0.002$ | $F_{1,11} = 0.09$, $p = 0.78$, $\eta^2_p = 0.01$ | $F_{1,11} < 0.01$, $p = 0.93$, $\eta^2_p = 0.0008$ |
| Allogrooming [%] | $F_{1,11} = 1.68$, $p = 0.22$, $\eta^2_p = 0.13$ | $F_{1,11} = 1.08$, $p = 0.32$, $\eta^2_p = 0.37$ | $F_{1,11} = 0.96$, $p = 0.35$, $\eta^2_p = 0.08$ |
| Total Social Behavior [%] | $F_{1,11} = 0.32$, $p = 0.58$, $\eta^2_p = 0.002$ | $F_{1,11} = 2.32$, $p = 0.16$, $\eta^2_p = 0.17$ | $F_{1,11} = 0.15$, $p = 0.71$, $\eta^2_p = 0.05$ |

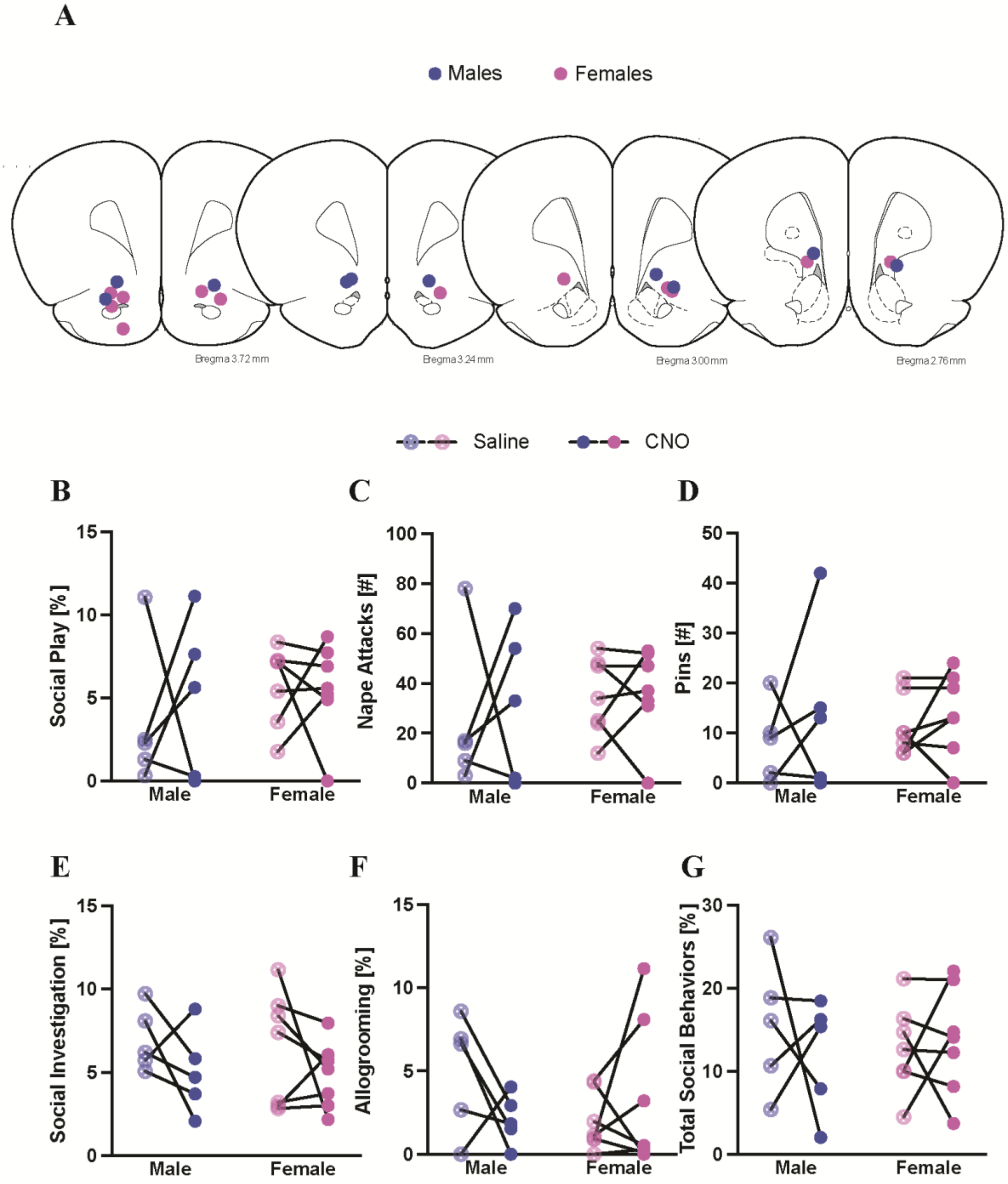


**Supplemental Figure 3. Local infusion of CNO outside of the NAc in hM3Dq transfected rats does not alter social play nor social investigation in juvenile males or females.** (A) Schematic depicting the location of cannula placements outside of the NAc. Both males and females showed similar duration of social play adapted from Paxinos and Watson (2007) (B), social investigation (E), allogrooming (F), and total social behaviors (G), as well as frequency of nape attacks (C) and pins (D), when treated with saline or CNO; repeated measures 2-way ANOVA, $p > 0.05$ for all.

**Supplemental Table 2.** Experiment 4: Effects of CNO infusion outside of the NAc in hM3Dq transfected rats; Two-way ANOVA statistics for behaviors quantified in social play testing. Significant main effects and interactions are notated in **bold**.

| | Drug Treatment | Sex | Drug x Sex |
|---|---|---|---|
| Social Play [%] | $F_{1,10} = 0.12$, $p = 0.73$, $\eta^2_p = 0.01$ | $F_{1,10} = 1.72$, $p = 0.22$, $\eta^2_p = 0.07$ | $F_{1,10} = 0.23$, $p = 0.64$, $\eta^2_p = 0.02$ |
| Nape Attacks [#] | $F_{1,10} = 0.15$, $p = 0.71$, $\eta^2_p = 0.01$ | $F_{1,10} = 0.77$, $p = 0.40$, $\eta^2_p = 0.04$ | $F_{1,10} = 0.07$, $p = 0.79$, $\eta^2_p = 0.01$ |
| Pins [#] | $F_{1,10} = 1.10$, $p = 0.32$, $\eta^2_p = 0.10$ | $F_{1,10} = 0.11$, $p = 0.75$, $\eta^2_p = 0.01$ | $F_{1,10} = 0.20$, $p = 0.67$, $\eta^2_p = 0.02$ |
| Social Investigation [%] | $F_{1,10} = 2.73$, $p = 0.13$, $\eta^2_p = 0.21$ | $F_{1,10} = 0.12$, $p = 0.74$, $\eta^2_p = 0.01$ | $F_{1,10} = 0.03$, $p = 0.87$, $\eta^2_p = 0.003$ |
| Allogrooming [%] | $F_{1,10} = 0.31$, $p = 0.59$, $\eta^2_p = 0.03$ | $F_{1,10} = 0.47$, $p = 0.51$, $\eta^2_p = 0.04$ | $F_{1,10} = 2.52$, $p = 0.14$, $\eta^2_p = 0.20$ |
| Total Social Behavior [%] | $F_{1,10} = 0.39$, $p = 0.54$, $\eta^2_p = 0.04$ | $F_{1,10} = 0.02$, $p = 0.91$, $\eta^2_p = 0.0007$ | $F_{1,10} = 0.23$, $p = 0.64$, $\eta^2_p = 0.02$ |

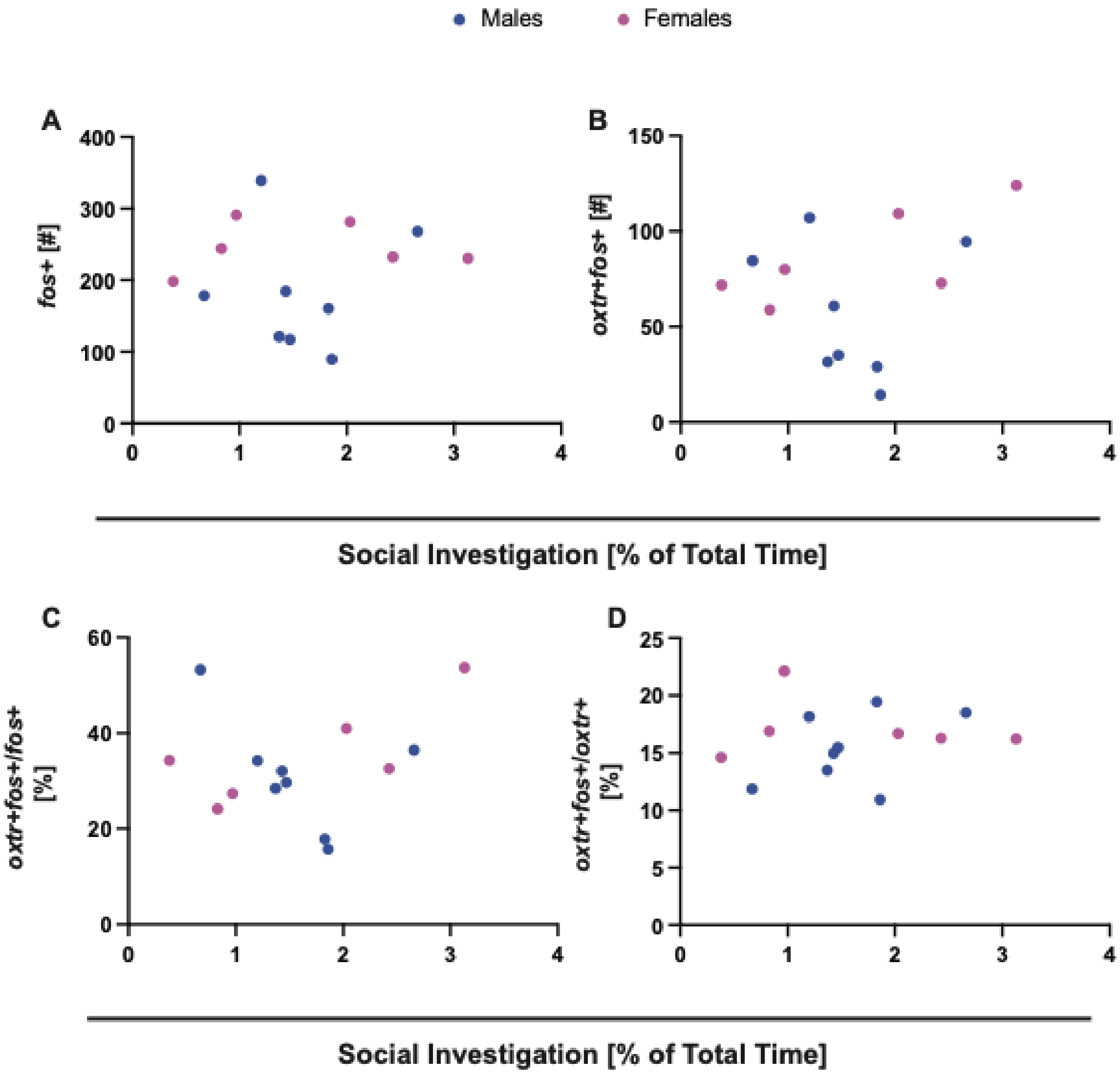


**Supplemental Figure 4. Social investigation does not correlate with the activation patterns within the anterior NAc** (A-D; Pearson's Correlation, $p > 0.05$ for all).
